# Regulation of Sex-biased Gene Expression by the Ancestral X-Y Chromosomal Gene Pair *Kdm5c-Kdm5d*

**DOI:** 10.1101/2024.10.24.620066

**Authors:** Rebecca M. Malcore, Milan Kumar Samanta, Sundeep Kalantry, Shigeki Iwase

## Abstract

Conventionally, Y-linked *Sry* is thought to drive sex differences by triggering differential hormone production. Ancestral X-Y gene pairs, however, are hypothesized to drive hormone-independent sex differences. Here, we show that the X-Y gene pair *Kdm5c*-*Kdm5d* regulates sex-biased gene expression in pluripotent mouse embryonic stem cells (ESCs). Wild-type (WT) XX female ESCs exhibit >2-fold higher expression of 409 genes relative to WT XY male ESCs. Conversely, WT XY male ESCs exhibit >2-fold higher expression of 126 genes compared to WT XX female ESCs. Loss of *Kdm5c* in female ESCs downregulates female-biased genes. In contrast, loss of either *Kdm5c* or *Kdm5d* in male ESCs upregulates female-biased genes and downregulates male-biased genes, effectively neutralizing sex-biased gene expression. KDM5C promotes the expression of *Kdm5d* and several other Y-linked genes in male ESCs. Moreover, ectopic *Kdm5d* expression in female ESCs is sufficient to drive male-biased gene expression. These results establish *Kdm5c*-*Kdm5d* as critical regulators of sex-biased gene expression.

## Introduction

Mammalian sexual differentiation is thought to be determined by the presence or absence of the Y chromosome. In therian mammals, males harbor one X and one Y chromosome, and females harbor two X chromosomes. During embryonic development, expression of the Y-linked *Sry* gene triggers differential gonad specification and sex hormone production in males (XY) relative to females (XX)^1^. Sex hormones drive sex differences in the reproductive tract as well as in non-reproductive tissues^2–4^.

*Sry* expression in mice begins between embryonic days (E) 10.5-11.5 and in humans between weeks ∼6-7 of gestation^1,5^. Prior to *Sry* expression, the sexes do not differ in sex hormone production or exposure. Despite the absence of *Sry* expression, however, early mammalian embryos exhibit sex differences. Early female mouse and human embryos are smaller and have reduced cell numbers relative to males^6–9^. Gene expression in early female vs. male embryos also differs^10–12^. These *Sry*-independent sex differences must arise from differences in the expression of X- or Y-encoded factors other than SRY. However, the X- or Y-linked genes responsible for these sex differences remain largely unknown.

The X and Y sex chromosomes are hypothesized to have originated from an equivalent pair of autosomes^13–15^. A series of inversions on the Y chromosome is believed to have suppressed crossing-over with the X chromosome^16^, which in turn led to substantial gene loss on the Y chromosome^14,15^. To compensate for Y chromosome gene loss, homologous X-linked genes became upregulated in males^13,17,18^. Similar upregulation occurred on both X chromosomes in females, resulting in excessive X-linked gene expression in females relative to males^13,17,18^. To reduce the expression of X-linked genes, females evolved X chromosome inactivation as a dosage compensation mechanism^13,19^. During X-inactivation, genes on one of the two X chromosomes are transcriptionally silenced in female cells^19^.

Given that X-inactivation largely equalizes X-linked gene expression between the sexes, Y-linked genes other than *Sry* are intuitive candidates for regulating sex differences since these genes are uniquely expressed in males. X-linked genes, though, may also contribute to sex differences^20^. Prior to the initiation of X-inactivation during early embryogenesis, X-linked genes are subject to higher expression in XX female cells relative to XY male cells^18,21,22^ and may also contribute to differences between early female and male embryos. Furthermore, once X-inactivation is established, a subset of X-linked genes is nevertheless capable of being more highly expressed in females relative to males by escaping X-inactivation^20,23^. Thus, both X- and Y-linked genes are candidate regulators of sex differences between females and males.

Several X-linked genes that constitutively escape X-inactivation have Y-linked homologs, forming X-Y gene pairs^14,15^. These X-Y gene pairs represent ancestral homologs in which the Y-linked copy has survived the decay of the therian Y chromosome^14,15^. Mice have nine X-Y gene pairs: *Ddx3x*-*Ddx3y*, *Eif2s3x*-*Eif2s3y*, *Kdm5c*-*Kdm5d*, *Rbmx/Rbmy*, *Sox3/Sry*, *Uba1*-*Ube1y1*, *Usp9x*-*Usp9y*, *Utx*-*Uty*, and *Zfx*-*Zfy1/2*. Four of these X-Y gene pairs are conserved across eutherian mammals, ubiquitously expressed, and harbor gene regulatory functions (*Ddx3x*-*Ddx3y*, *Eif2s3x*-*Eif2s3y*, *Kdm5c*-*Kdm5d*, and *Utx*-*Uty*)^14,15^. If both the function and expression output of a particular X-Y gene pair is equivalent between females and males, then the X-Y homologs would be analogous to biallelically expressed autosomal genes^15^. However, growing evidence suggests that homologous X-Y gene pairs have diverged during evolution and have distinct functions^24–28^. Divergent functions of X-Y gene pairs, in turn, may shape sex hormone-independent sex differences^24^.

Among the conserved X-Y gene pairs, the ancestral X-linked *Kdm5c* and Y-linked *Kdm5d* gene pair has emerged as candidate regulators of sex differences. *Kdm5c* and *Kdm5d* both encode ubiquitously expressed lysine demethylases that remove histone H3 lysine 4 di- and tri-methylation modifications (H3K4me2/3), which are associated with transcriptionally engaged promoters and enhancers^29–31^. Several studies have implicated KDM5C in sex differences of adult tissues, including adipose tissues, bone, and the brain^32–34^. However, these studies did not address the roles of KDM5D, which may or may not compensate for the absence of KDM5C in males. Reduction in KDM5D by siRNA knockdown has been implicated in the regulation of sex-biased gene expression in mouse embryonic fibroblast (MEF) cells^35^. KDM5C, however, was not tested in this study^35^. Thus, the similar vs. divergent functions of KDM5C vs. KDM5D in regulating sex-differences remain unknown.

In this study, we systematically investigated the gene regulatory roles of *Kdm5c* and *Kdm5d* in mouse embryonic stem cells (ESCs). ESCs represent a developmental stage prior to the expression of *Sry*, i.e. prior to the onset of sex hormone differences between the sexes. We find overlapping and divergent roles of *Kdm5c* and *Kdm5d* in driving sex-biased gene expression in mouse ESCs.

## Results

### Sex-biased gene expression in wild-type female and wild-type male ESCs

To define sex-biased genes in mouse ESCs, we generated and compared female (*X*^5c–fl^*X*^5c–fl^) and male (*X*^5c–fl^*Y*) ESC lines with intact *Kdm5c* and *Kdm5d* alleles, herein referred to as wild-type (WT) female and WT male ESCs. We verified chromosomal copy number integrity in these lines through low-pass whole genome sequencing (WGS) (Figure S1A-B). We also tested the pluripotent state of the WT ESC lines by analyzing expression of pluripotency-associated genes by poly(A)-enriched RNA sequencing (RNA-seq) and found comparable expression in the ESC lines (Kruskal-wallis test p = 0.52; Figure S1D). We then performed a genome-wide differential gene expression analysis in these WT ESC lines (Figure 1A). Genes were designated as differentially expressed if they were >2-fold (Log_2_ of 1) higher in one sex relative to the other and had a false discovery rate (FDR) of <0.01. Of the 12,627 genes expressed in both female and male ESCs (≥ 1 count per million; CPM), 535 genes (4.2%) were differentially expressed between females and males (Figure 1B). Of these, 409 genes were more highly expressed in WT female compared to WT male ESCs (‘female-biased genes’; Figure 1A), and 126 genes were more highly expressed in WT male compared to WT female ESCs (‘male-biased genes’; Figure 1A).

**Figure 1:**
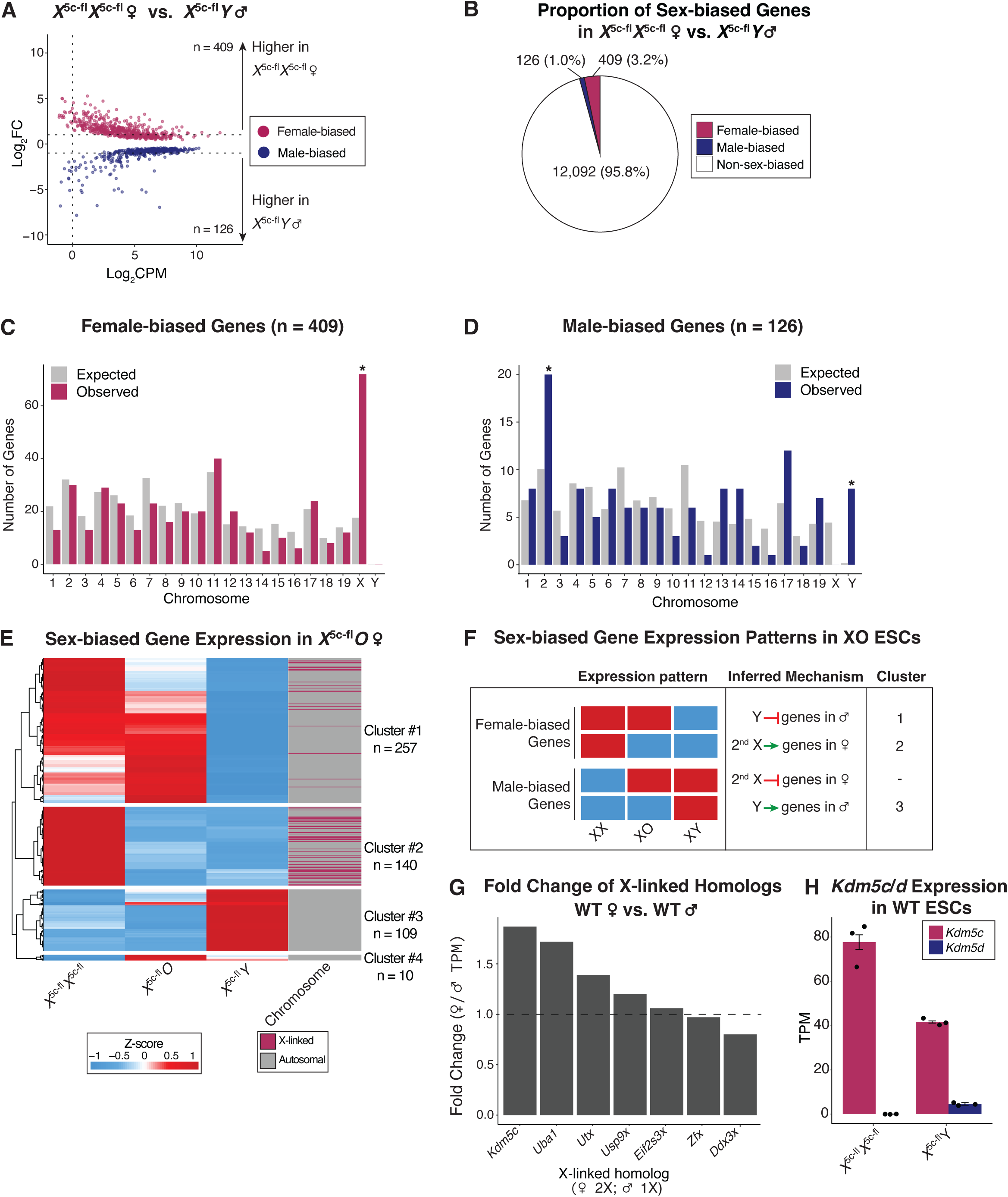
Sex-biased gene expression in WT *X*^5c-fl^*X*^5c-fl^ female and WT *X*^5c-fl^*Y* male ESCs. (A) RNA-seq MA plot comparing WT *X*^5c-fl^*X*^5c-fl^ female vs. WT *X*^5c-fl^*Y* male ESCs (3 biological replicates each). ‘n’; number of differentially expressed genes (≥ 1 Log_2_ fold change (Log_2_FC), ≥ 1 count per million (CPM), false discovery rate (FDR) < 0.01). Maroon, female-biased genes. Dark blue, male-biased genes. Grey, non-sex-biased genes. Only genes with an FDR < 0.01 were plotted. (B) Proportion of expressed genes (CPM ≥ 1) in WT *X*^5c-fl^*X*^5c-fl^ female and WT *X*^5c-fl^*Y* male ESCs that exhibit sex-biased expression. (C and D) Distribution of female-biased (C) or male-biased (D) genes on each chromosome. Expected values were determined by Chi-square test. (*); p < 0.05 (binomial test, Benjamini-Hotchberg corrected) for difference between expected and observed number of sex-biased genes. (E) Heatmap of sex-biased gene expression (TPM) in XX, XO, and XY ESCs. Heatmap is clustered and scaled by row and colored by Z-score. Cluster cut-offs were determined by tree height (> 2). (F) Expression patterns of sex-biased genes in XX, XO, and XY ESCs and the inferred mechanisms by which the X and the Y chromosomes influence sex-biased gene expression. Cluster numbers correspond to (E). (G) Relative X-linked homolog expression (TPM) in WT *X*^5c-fl^*X*^5c-fl^ female vs. WT *X*^5c-fl^*Y* male ESCs. (H) *Kdm5c* and *Kdm5d* RNA expression in the WT *X*^5c-fl^*X*^5c-fl^ female vs. WT *X*^5c-fl^*Y* male ESCs (TPM; mean ± SEM).

X-linked genes were enriched among female-biased genes (binomial test p = 6.9E-22; Figure 1C). Conversely, no X-linked genes were male-biased (Figure 1D). The enrichment of X-linked genes among female-biased genes is consistent with the presence of two transcriptionally active X chromosomes in female ESCs. Y-linked genes were enriched in male-biased genes (binomial test p = 1.38E-08), as expected, as well as chromosome 2 genes (binomial test p = 0.041; Figure 1D).

We next sought to determine the impact of the second X chromosome in females and the Y chromosome in males on autosomal sex-biased gene expression (Figure 1E and F). To this end, we employed XO female ESCs (*X*^5c–fl^*O*; Methods), which lack a second X chromosome, and compared sex-biased gene expression across WT XX, XY, and XO ESCs. From these comparisons, we inferred the contribution of the second X chromosome and the Y chromosome to sex-biased gene expression (Figure 1F). By hierarchical clustering, sex-biased genes clustered into four groups with distinct expression patterns in the XX, XO, and XY ESCs (Figure 1E and F).

Female-biased genes separated into two clusters (Clusters 1 and 2; Figure 1E and F). Genes in Cluster 1 were upregulated in both XX and XO females relative to XY males. This pattern is consistent with the presence of the Y chromosome repressing Cluster 1 genes in XY cells, thereby resulting in their female expression bias. Cluster 2 female-biased genes were downregulated in both XO and XY ESCs relative to XX ESCs. Cluster 2 contained a high proportion of X-linked genes (n = 46 of 140), unlike Cluster 1 (Figure 1E). This pattern is consistent with the second active X chromosome upregulating Cluster 2 genes in XX ESCs relative to XO and XY ESCs. Together, this analysis suggests that female-biased gene expression is achieved by both the absence of the Y chromosome and the presence of the second X chromosome.

The expression of male-biased genes also separated into two clusters (Clusters 3 and 4; Figure 1E and F). Cluster 3 contained almost all male-biased genes (n = 109), which were upregulated in XY ESCs relative to both XX and XO ESCs (Figure 1E). This pattern is consistent with the Y chromosome upregulating the expression of most male-biased genes.

The remaining male-biased genes separated into Cluster 4 (n = 10), which unexpectedly exhibited the highest expression in XO females (Figure 1E). Cluster 4 genes are potentially repressed by both the Y chromosome in XY males and the second X chromosome in XX females. Interestingly, the hierarchical clustering did not yield a cluster of male-biased genes that are suppressed by the second X chromosome in XX ESCs (Figure 1F), suggesting that the X chromosome does not carry suppressors of male-biased genes. Taken together, these results indicate that both the Y chromosome and the second X chromosome shape sex-biased gene expression in ESCs. Moreover, the Y chromosome exerts a greater influence on sex-biased gene expression relative to the second X chromosome by both promoting male-biased gene expression and repressing female-biased gene expression.

To define candidate X- and Y-linked regulators of sex-biased gene expression in ESCs, we first considered the single-copy Y-linked genes expressed in WT male ESCs (≥ 1 CPM, n = 8; Figure 1D). Notably, all expressed single-copy Y-linked genes in WT male ESCs have X-linked homologs. Among the expressed X-linked homologs of X-Y gene pairs, *Kdm5c* exhibits the greatest fold change in expression between females vs. males (1.9x; Figure 1G and H). Moreover, in WT male ESCs, *Kdm5c* expression is 9x higher than its Y-linked homolog *Kdm5d* (Figure 1H). *Kdm5c* and *Kdm5d* encode H3K4me2/3 demethylases and are thus positioned to function as transcriptional regulators of sex-biased gene expression^29–31,36^.

### Altered sex-biased gene expression in *Kdm5c-* and *Kdm5d*-mutant ESCs

To compare the gene regulatory functions of KDM5C and KDM5D in female and male ESCs, we derived a series of mutant *Kdm5c* and *Kdm5d* ESC lines using conditional alleles as shown in Figure 2. The conditional *Kdm5c* allele harbors *loxP* sequences flanking exons 11 and 12, which encode the JumonjiC (JmjC) demethylase domain (*X*^5c–fl^; Figure 2A)^37^. We also generated mice harboring a conditional *Kdm5d* allele, where exons 11 and 12 encoding the JmjC domain were again flanked by *loxP* sequences (*Y*^5d–fl^; Figure 2 and Figure S2). We bred mice harboring the *X*^5c–fl^ or *Y*^5d–fl^ alleles with a germline *Cre* line to obtain *Kdm5c-* or *Kdm5d*-mutant alleles (*X*^5c–Δ^ and *Y*^5d–Δ^). From the resultant embryos, we derived all combinations of *Kdm5c*- and *Kdm5d*-mutant ESC lines (Figure 2C). The *X*^5c–fl^*X*^5c–fl^ female and *X*^5c–fl^*Y* male ESCs analyzed in Figure 1 serve as WT controls. *Kdm5c*-heterozygous (*X*^5c–fl^*X*^5c–Δ^) and *Kdm5c*-homozygous (*X*^5c–Δ^*X*^5c–Δ^) ESC lines comprise the mutant female genotypes. *Kdm5c*-hemizygous (*X*^5c–Δ^*Y*), *Kdm5d*-hemizygous (*XY*^5d–Δ^), and *Kdm5c/d* double-mutant (*X*^5c–Δ^*Y*^5d–Δ^) ESC lines are the mutant male genotypes. Low-pass whole genome sequencing validated the integrity of all autosomes in the mutant ESC lines (Figure S3A). We also confirmed loss of the JmjC-encoding exons in *Kdm5c-* (Figure 2D) and *Kdm5d-*mutant (Figure 2E) genotypes by RNA-seq. We found comparable expression of pluripotency genes in the mutant ESC lines relative to the WT ESC lines (Kruskal-wallis test p = 0.97; Figure S3B).

**Figure 2:**
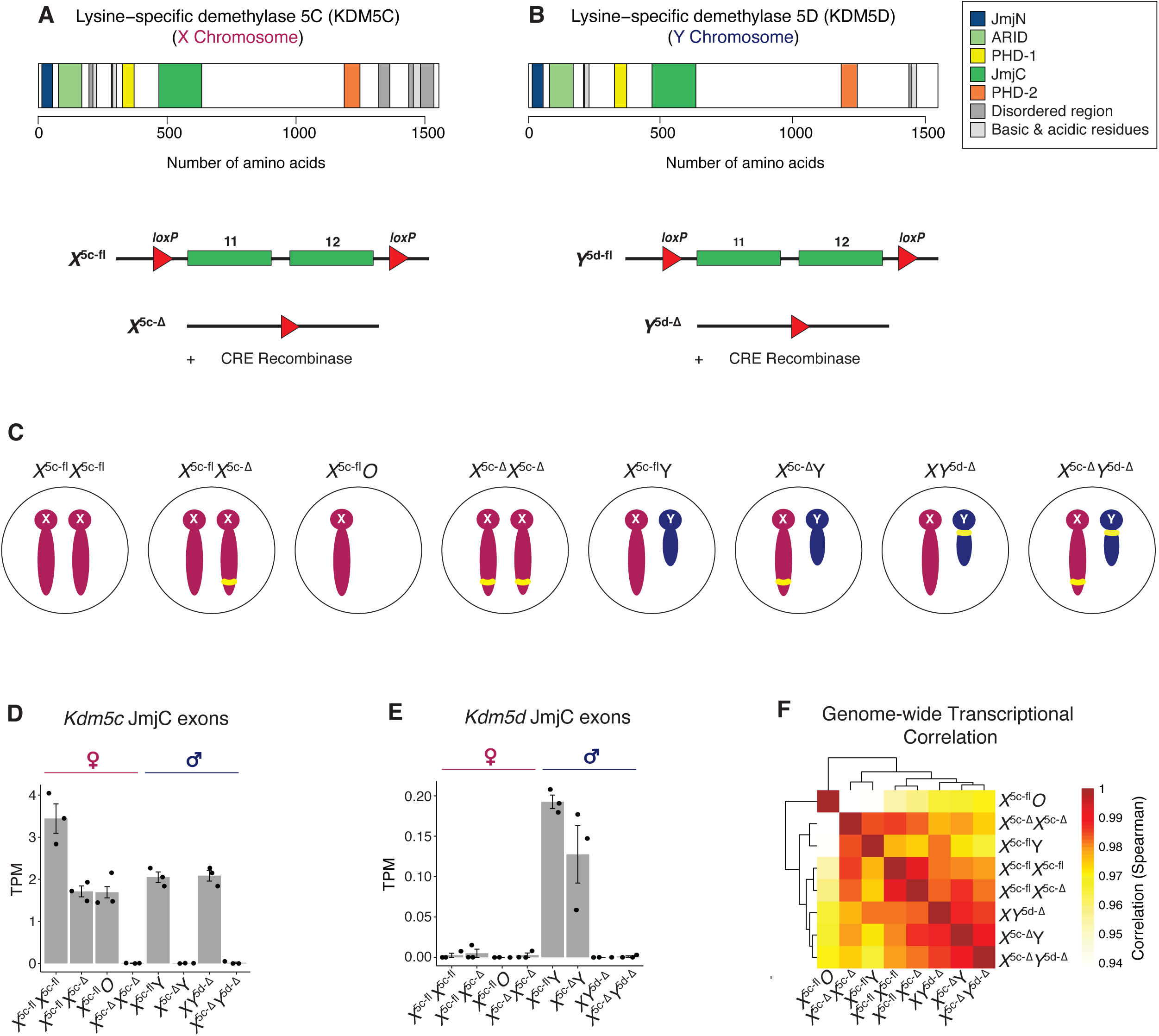
Generation of *Kdm5c* and *Kdm5d* floxed and mutant ESCs. (A and B) Conditional mutation strategy of *Kdm5c* (A; ref. 37) and *Kdm5d* (B; this study). In both cases, exons encoding the enzymatic JmjC domain were floxed (fl) and conditionally deleted (Δ) via germline *Cre* recombinase. (C) Schematic of all *Kdm5c* and *Kdm5d* WT and mutant male and female ESC lines used in this study. Yellow bar, *Kdm5c* mutation on the X chromosome or *Kdm5d* mutation on the Y chromosome. (D and E) Expression (TPM) of *Kdm5c* (D) or *Kdm5d* (E) JmjC targeted exons in the WT and mutant ESCs (mean ± SEM; 3 biological replicates each). (F) Spearman correlations of the transcriptomes (TPM) of all WT and mutant ESCs. Heatmap is scaled by row and colored by Z-score.

We initially compared global gene expression across all control and mutant ESC genotypes. Overall, all genotypes exhibited high similarity in gene expression (Spearman correlation ≥ 0.94; Figure 2F). However, the *X*^5c–Δ^*Y*, *XY*^5d–Δ^, and *X*^5c–Δ^*Y*^5d–Δ^ mutant male ESC transcriptomes were more similar to that of WT female ESCs compared to WT male ESCs (Spearman correlation 0.98; Figure 2F). Conversely, gene expression in homozygous *X*^5c–Δ^*X*^5c–Δ^ mutant female ESCs was more similar to WT male ESCs compared WT female ESCs (Spearman correlation 0.98; Figure 2F). These observations suggest that female- and male-biased gene expression are altered in the absence of KDM5D and/or KDM5C. The expression levels of sex hormone nuclear receptor genes encoding the androgen (*Ar*), estrogen (*Esr1* and *Esr2*) and progesterone (*Pgr*) receptors were very low (<0.4 transcripts per million; TPM) in all ESC lines (Table S1), and all cell lines were cultured in the same conditions. Thus, any changes in sex-biased gene expression from the loss of *Kdm5c* or *Kdm5d* are likely independent of sex hormones.

### Sex-biased gene expression is neutralized in *X*^5c–Δ^*Y*, *XY*^5d–Δ^, and *X*^5c–Δ^*Y*^5d–Δ^ male ESCs

To define the changes in sex-biased gene expression in *Kdm5c-* and *Kdm5d*-mutant ESCs, we first tested the expression of sex-biased genes as defined in WT female vs. WT male ESCs (Figure 1) in the *X*^5c–Δ^*Y*, *XY*^5d–Δ^, and *X*^5c–Δ^*Y*^5d–Δ^ mutant male ESCs (Figure 3A). Sex-biased gene expression in the mutant male ESCs did not resemble that of WT male ESCs, and instead the mutants exhibited upregulation of female-biased genes and downregulation of male-biased genes (Figure 3A). *X*^5c–Δ^*Y* ESCs upregulated 171 female-biased genes and downregulated 70 male-biased genes (Figure 3B). *XY*^5d–Δ^ ESCs upregulated 88 female-biased genes and downregulated 23 male-biased genes (Figure 3C). *X*^5c–Δ^*Y*^5d–Δ^ ESCs upregulated 200 female-biased genes and downregulated 75 male-biased genes (Figure 3D). Notably, no male-biased genes were upregulated and no female-biased genes were downregulated in any of the mutant male genotypes relative to WT male ESCs. All three mutant male genotypes also showed dysregulation of non-sex biased genes (Figure 3B-D, Figure S4A). However, sex-biased genes were significantly enriched amongst all differentially expressed genes in *X*^5c–Δ^*Y* (logistic regression p = 5.6e-22), *XY*^5d–Δ^ (p = 1.9e-06), and *X*^5c–Δ^*Y*^5d–Δ^ (p = 1.0e-28) male ESCs relative to WT *X*^5c–fl^*Y* male ESCs (Figure 3E). As a result, all mutant male genotypes expressed fewer female- and male-biased genes when compared to WT female ESCs (Figure 3F). Together, these analyses suggest that both KDM5C and KDM5D repress female-biased genes and activate male-biased genes in male ESCs.

**Figure 3:**
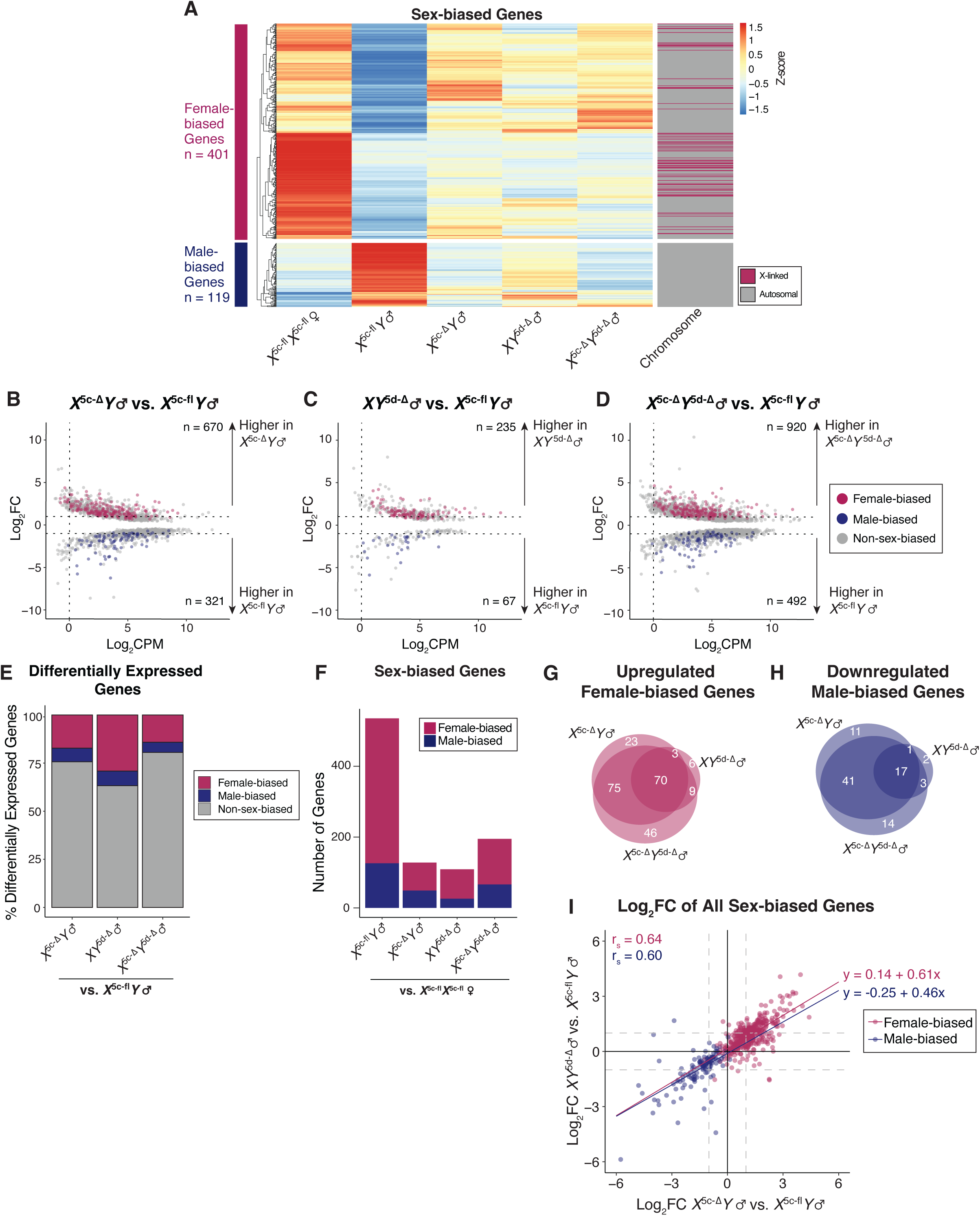
Neutralization of sex-biased gene expression in *Kdm5c*- and *Kdm5d*-mutant male ESCs. (A) Heatmap of sex-biased gene expression (TPM) in WT *X*^5c-fl^*X*^5c-fl^ female and WT *X*^5c-fl^*Y* male ESCs vs. *X*^5c-Δ^*Y*, *XY*^5d-Δ^, and *X*^5c-Δ^ *Y*^5d-Δ^ mutant male ESCs. Heatmap is clustered by row, scaled by row, and colored by Z-score. Cluster cut-offs were determined by tree height (≥ 3.5). (B-D) RNA-seq MA plots of pairwise comparisons of *X*^5c-Δ^*Y* (B), *XY*^5d-Δ^ (C), and *X*^5c-Δ^ *Y*^5d-Δ^ (D) mutant male ESCs vs. WT *X*^5c-fl^*Y* male ESCs. Maroon, female-biased genes. Dark blue, male-biased genes. Grey, non-sex-biased genes. Only genes with an FDR < 0.01 were plotted. (E) The proportions of differentially expressed genes in *X*^5c-Δ^*Y*, *XY*^5d-Δ^, and *X*^5c-Δ^ *Y*^5d-Δ^ mutant male ESCs that exhibit sex-biased expression. (F) Number of sex-biased genes determined by pairwise comparisons between WT *X*^5c-fl^*X*^5c-fl^ female ESCs vs. *X*^5c-Δ^*Y*, *XY*^5d-Δ^, and *X*^5c-Δ^ *Y*^5d-Δ^ mutant male ESCs. (G-H) Overlap of differentially expressed female-biased genes (G) and male-biased genes (H) in *X*^5c-Δ^*Y*, *XY*^5d-Δ^, and *X*^5c-Δ^ *Y*^5d-Δ^ mutant male ESCs. (I) Correlation of sex-biased gene expression Log_2_FCs in *X*^5c-Δ^*Y* and *XY*^5d-Δ^ mutant male ESCs. Maroon, female-biased genes. Dark blue, male-biased genes. Linear models were determined by y ∼ x. r_s_, Spearman correlation. Three biological replicates were used per genotype and all ‘n’ numbers indicate genes that were filtered for ≥ 1 Log_2_FC, ≥ 1 CPM, and FDR < 0.01.

We next examined the relative contributions of KDM5C and KDM5D in regulating gene expression in male ESCs. Compared to *XY* ^5d–Δ^ ESCs, *X*^5c–Δ^*Y* ESCs displayed a greater number of upregulated and downregulated genes genome-wide (Figure S4A). A subset of genes, however, were uniquely dysregulated in *XY* ^5d–Δ^ ESCs but not *X*^5c–Δ^*Y* ESCs (Figure S4A). A similar pattern was observed when comparing the dysregulated female- and male-biased genes in *X*^5c–Δ^*Y* and *XY* ^5d–Δ^ ESCs (Figure 3G and H). Almost all female- and male-biased genes exhibited similar directional changes in expression in both the *X*^5c–Δ^*Y* ESCs and *XY* ^5d–Δ^ ESCs despite not all reaching statistical significance in differential expression (Log_2_FC relative to WT male ESCs, r_s_ = 0.64 and 0.60; Figure 3I). Linear regression analysis, though, demonstrates that KDM5C has a stronger impact on sex-biased gene expression compared to KDM5D (slope = 0.61 and 0.46; Figure 3I). Examples of sex-biased genes whose expression changes similarly in the absence of *Kdm5c* and *Kdm5d* include female-biased *Tspo2* and male-biased *Slc44a2* (Figure S4B and C). Examples of sex-biased genes that change in expression only in the absence of *Kdm5c* include female-biased *Xlr3a* and male-biased *Creb1*. And, examples of sex-biased genes that change in expression only in the absence of *Kdm5d* include female-biased *Egln3* and male-biased *Tcerg1l* (Figure S4D-G).

The presence of the second X chromosome in female ESCs upregulates the expression of a subset of autosomal female-biased genes (Cluster 2; Figure 1E and F). We thus sought to test whether upregulation of X-linked genes coincides with the increased expression of female-biased genes in *Kdm5c-* and *Kdm5d-*mutant male ESCs. Bulk X-chromosomal gene expression was similar between *Kdm5c-* and *Kdm5d-*mutant male ESCs and both ESC genotypes were similar to WT male ESCs and did not resemble WT female ESCs (Figure S5A). Among the differentially expressed X-linked genes, the three mutant male genotypes did not share any upregulated X-linked genes (Figure S5B). These results suggest that upregulation of X-linked genes in *Kdm5c*- and *Kdm5d*-mutant male ESCs does not explain their upregulation of autosomal female-biased genes.

The presence of the Y chromosome represses a subset of autosomal female-biased genes and upregulates almost all autosomal male-biased genes (Clusters 1 and 3 in Figure 1E and F). Thus, reduced Y-linked gene expression in *Kdm5c*- or *Kdm5d*-mutant male ESCs could drive changes in both female- and male-biased gene expression in the mutant male ESCs. The majority of expressed single-copy Y-linked genes in male ESCs were downregulated in both the *X*^5c–Δ^*Y* and *X*^5c–Δ^*Y*^5d–Δ^ mutant ESCs (5 of 8: *Ddx3y*, *Eif2s3y*, *Usp9y*, *Uty*, and *Zfy2*; Figure S5C). Moreover, *Kdm5d* was downregulated in *X*^5c–Δ^*Y* and *X*^5c–Δ^*Y*^5d–Δ^ mutant ESCs (Figure S5C). Note that in *XY* ^5d–Δ^ ESCs, *Kdm5d* expression is unchanged because most reads originate from the intact exons (Figure 2E, Figure S5C). In *XY* ^5d–Δ^ ESCs Y-linked gene expression remained comparable to that of WT male ESCs, which is consistent with fewer sex-biased gene expression changes in these cells relative to the *X*^5c–Δ^*Y* and *X*^5c–Δ^*Y*^5d–Δ^ ESCs (Figure 3G and H, Figure S5C). Together, these results suggest that 1) *Kdm5c* promotes *Kdm5d* expression, and 2) the downregulation of several Y-linked genes may contribute to the increased upregulation of autosomal female-biased genes and downregulation of autosomal male-biased genes in *X*^5c–Δ^*Y* and *X*^5c–Δ^*Y*^5d–Δ^ ESCs relative to *XY* ^5d–Δ^ ESCs.

### Reduced sex-biased gene expression in *Kdm5c*-mutant female ESCs

To define changes in sex-biased gene expression in *Kdm5c*-mutant females, we next examined both *X*^5c–fl^*X*^5c–Δ^ heterozygous-mutant and *X*^5c–Δ^*X*^5c–Δ^ homozygous-mutant female ESCs (Figure 4A). We found that homozygous *X*^5c–Δ^*X*^5c–Δ^ ESCs exhibited more severely disrupted sex-biased gene expression relative to heterozygous *X*^5c–fl^*X*^5c–Δ^ ESCs (Figure 4A). Whereas homozygous *X*^5c–Δ^*X*^5c–Δ^ ESCs upregulated 8 male-biased genes and downregulated 54 female-biased genes, heterozygous *X*^5c–fl^*X*^5c–Δ^ ESCs upregulated 3 male-biased genes and downregulated 2 female-biased genes (Figure 4B and C).

**Figure 4:**
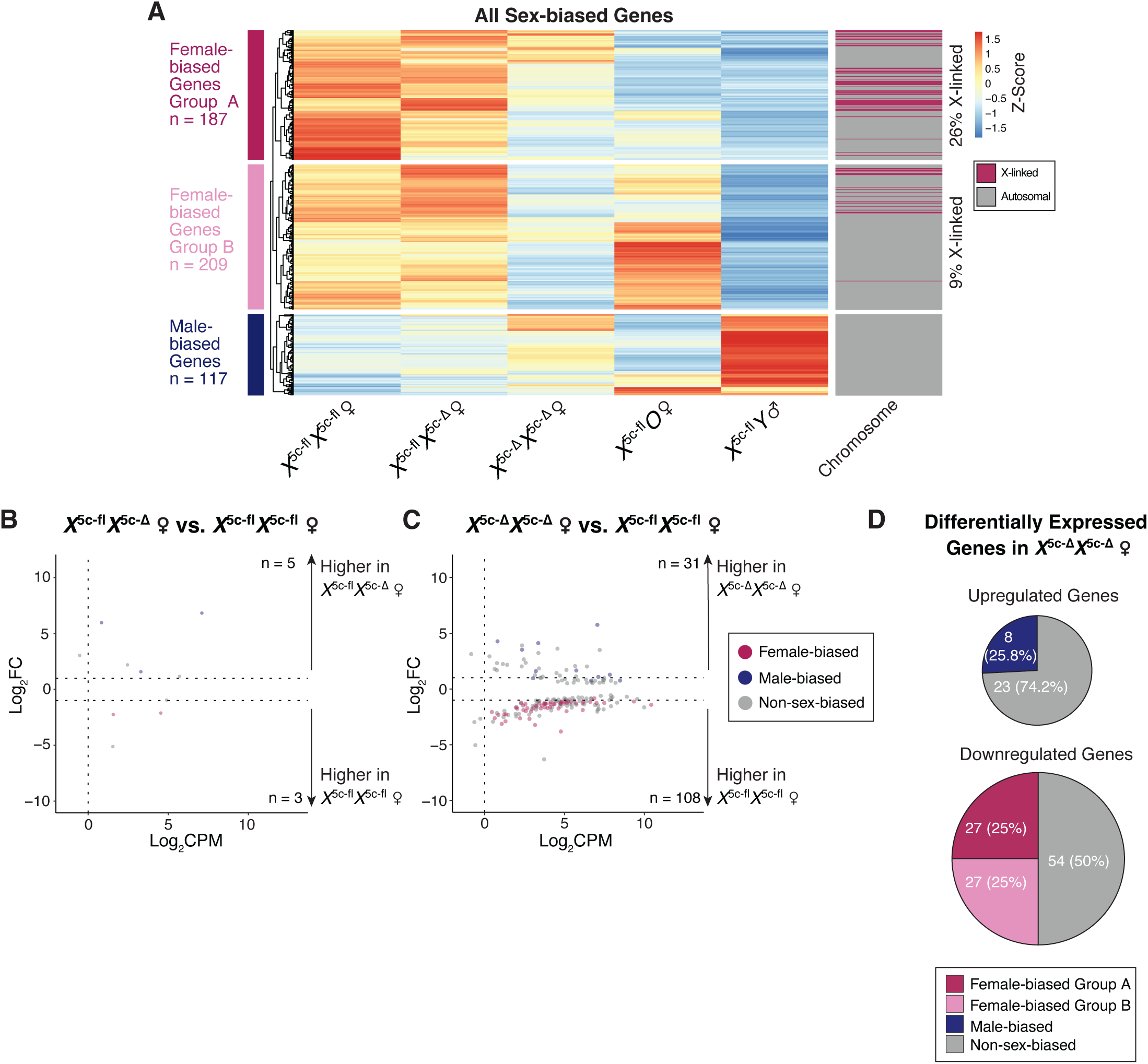
Reduced sex-biased gene expression in *Kdm5c*-mutant female ESCs. (A) Heatmap of sex-biased gene expression (TPM) in WT *X*^5c-fl^*X*^5c-fl^ female and WT *X*^5c-fl^*Y* male vs. *Kdm5c-*mutant female ESCs. Heatmap is clustered by row, scaled by row, and colored by Z-score. Cluster cut-offs were determined by tree height (≥ 3). (B-C) RNA-seq MA plots of pairwise comparisons of *X*^5c-fl^*X*^5c-Δ^ heterozygous mutant female (B) and *X*^5c-Δ^*X*^5c-Δ^ homozygous mutant female ESCs (C) vs. WT *X*^5c-fl^*X*^5c-fl^ female ESCs. Maroon; female-biased genes. Dark blue; male-biased genes. Grey; non-sex-biased genes. Only genes with an FDR < 0.01 were plotted. (D) Proportion of upregulated or downregulated genes in *X*^5c-Δ^*X*^5c-Δ^ homozygous mutant female ESCs that are sex-biased in expression. Three biological replicates were used per genotype and all ‘n’ numbers indicate genes that were filtered for ≥ 1 Log_2_FC, ≥ 1 CPM, and FDR < 0.01.

Female ESCs often lose one X chromosome in culture^38^. Homozygous *X*^5c–Δ^*X*^5c–Δ^ ESCs lost an X chromosome more rapidly compared to both the WT and heterozygous *X*^5c–fl^*X*^5c–Δ^ female ESCs (Figure S6). At the time of analysis (passage 9), homozygous *X*^5c–Δ^*X*^5c–Δ^ ESCs were a mosaic population with ∼30% of the cells harboring two X chromosomes and 70% with a single X chromosome (Figure S6). Changes in sex-biased gene expression in homozygous *X*^5c–Δ^*X*^5c–Δ^ ESCs may, therefore, be explained by X-monosomy in a large subset of the cells.

To determine the impact of X-monosomy on sex-biased gene expression changes in the *Kdm5c*-deficient female ESCs, we first compared gene expression changes in *X*^5c–Δ^*X*^5c–Δ^/*X*^5c–Δ^*O* ESCs to *X*^5c–fl^*O* ESCs. Hierarchical clustering of RNA-seq for all sex-biased genes revealed two distinct groups of female-biased genes (Figure 4A). Group A genes were downregulated in both *X*^5c–fl^*O* and the mosaic *X*^5c–Δ^*X*^5c–Δ^/*X*^5c–Δ^*O* ESCs relative to WT female ESCs, and thus were likely driven by X-monosomy (n = 187, Figure 4A). In agreement, 26% of group A female-biased genes were X-linked (n = 49 of 187, Figure 4A). Conversely, Group B female-biased genes were highly expressed in *X*^5c–fl^*O* ESCs and became downregulated in the mosaic *X*^5c–Δ^*X*^5c–Δ^/*X*^5c–Δ^*O* ESCs (n = 209; Figure 4A). This pattern suggests that Group B sex-biased expression changes are not influenced by X-monosomy. These data indicate that KDM5C maintains female-biased gene expression by both transcriptional regulation and X-chromosome retention.

Downregulated genes in the mosaic *X*^5c–Δ^*X*^5c–Δ^/*X*^5c–Δ^*O* ESCs were enriched for female-biased genes, with 27 Group A genes and 27 Group B genes significantly downregulated (logistic regression p = 2.0e-05; Figure 4C and D). However, only 8 male-biased genes were upregulated in the mosaic *X*^5c–Δ^*X*^5c–Δ^/*X*^5c–Δ^*O* ESCs, comprising 6.3% of the male-biased genes in WT male ESCs (Figure 1A, Figure 4D). Thus, KDM5C upregulates a subset of female-biased genes but has a limited role in repressing male-biased genes in female ESCs. These observations are consistent with the requirement of the Y chromosome for robust upregulation of the majority of male-biased genes in WT ESCs (Figure 1E and F).

### *Kdm5d* induces male-biased gene expression in XO female ESCs

The data thus far indicate that in ESCs, *Kdm5c* and *Kdm5d* are both necessary for sex-biased gene expression. We next tested whether Y-linked *Kdm5d* is sufficient to drive male-biased gene expression. Initially, we attempted to introduce a *Kdm5d* transgene via lentiviral. transduction in heterozygous *X*^5c–fl^*X*^5c–Δ^ female ESCs that harbor a single intact *Kdm5c* allele. However, all *Kdm5d*-transgene expressing clones had lost one X chromosome and were *X*^5c–fl^*O* (Figure 5A). These cells, however, still allowed us to examine the roles of *Kdm5d* in the absence of other Y-linked genes. The *X*^5c–fl^*O* transgenic Clones 1 and 2, which expressed the *Kdm5d* transgene below the levels of WT male ESCs, did not exhibit major alterations in sex-biased gene expression (Figure 5D). Strikingly, *X*^5c–fl^*O* transgenic Clone 3, expressing *Kdm5d* at comparable levels to WT male ESCs, exhibited downregulation of the majority of female-biased genes and upregulation of a subset of male-biased genes, resulting in a WT male-like transcriptome (Figure 5C and D). A subset of male-biased genes is not upregulated in Clone 3, suggesting that other Y-linked gene(s) are required to upregulate the remaining male-biased genes (Figure 5D). These results demonstrate that *Kdm5d* is sufficient to drive male-biased gene expression in cells lacking a Y chromosome.

**Figure 5:**
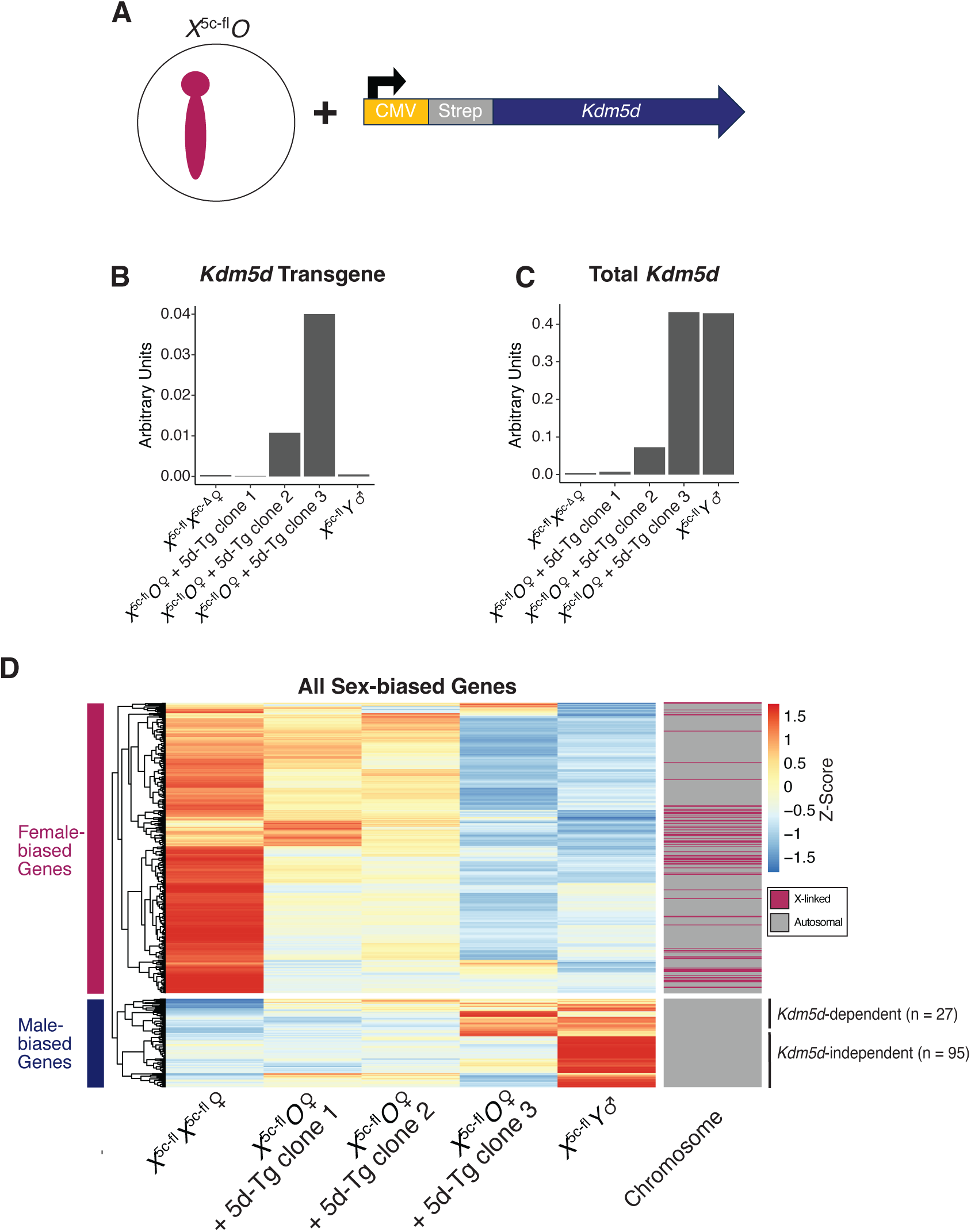
K*d*m5d is sufficient to induce male-biased gene expression in *X*^5c-fl^*O* ESCs. (A) Schematic depicting experimental approach. A *Kdm5d* transgene tagged at the N-terminus with a Strep tag and constitutively driven by a CMV promoter was introduced into *X*^5c-fl^*O* ESC clones by lentivirus. (B) The RNA expression levels of the *Kdm5d* transgene determined by RT-qPCR with a forward primer specific to the Strep tag and a reverse primer in the 5’ end of *Kdm5d*. Data are normalized to *Tbp* RNA. (C) The RNA expression levels of total *Kdm5d* by RT-qPCR with primers specific to *Kdm5d* exons 11 and 12. Data are normalized to *Tbp* RNA. (D) Heatmap of sex-biased gene expression (TPM) in WT *X*^5c-fl^*X*^5c-fl^ female and WT *X*^5c-fl^*Y* male ESCs vs. *X*^5c-fl^*O Kdm5d*-transgenic clone ESCs. Heatmap is clustered by row, scaled by row, and colored by Z-score. Cluster cut-offs were determined by tree height (≥ 4).

## Discussion

The X and Y chromosomes are the sources of differences between the mammalian sexes. The sex chromosomes are hypothesized to have originated from a homologous pair of autosomes that differentiated during evolution^13–15^. X-Y differentiation led to the loss of a vast majority of genes on the Y chromosome with homologs on the X chromosome^14,15^. Notably, a subset of ancestral homologous genes is retained on the X and Y chromosomes, comprising X-Y gene pairs^14,15^. A number of these X-Y gene pairs are ubiquitously expressed and harbor regulatory functions^15^. X-Y gene pairs expressed equivalently between the sexes are expected to function similarly, akin to two alleles of an autosomal gene^15^. However, growing evidence suggests that many X-Y gene pairs have diverged in function due to the accumulation of sequence differences during evolution^24–28^. This sequence divergence can lead to differences in the level of expression or encoded amino acid sequence that results in dissimilar biochemical activities of the X and Y homologs. Through these mechanisms, X-Y gene pairs may thus drive differences between the sexes^24^. Five X-Y gene pairs are conserved across eutherians and are positioned to regulate sex differences: *Sox3*/*Sry, Utx*/*Uty*, *Zfx*/*Zfy, Ddx3x*/*Ddx3y*, and *Kdm5c*/*Kdm5d*^14,15^.

The Y-linked *Sry* gene is thought to the primary driver of sex differences. SRY induces testis differentiation, which in turn produces the hormone testosterone required for male sexual differentiation^1^. *Sry* has a diverged X-linked homolog, *Sox3*, which is not required for male sexual differentiation^39^. When overexpressed, however, SOX3 can cause testis differentiation in XX female mice^39^, suggesting overlapping biochemical functions encoded by the *Sox3*-*Sry* X-Y gene pair.

Recent findings suggest that other conserved eutherian X-Y gene pairs may also contribute to sex differences. The X-linked histone H3K27me3 demethylase *Utx* regulates autosomal sex-biased gene expression in male mouse ESCs^26^. *Utx* mutant male ESCs exhibit a loss of 27-32% of sex-biased gene expression relative to WT ESCs. However, whether Y-linked *Uty* functions similarly to *Utx* and whether *Utx* functions similarly in female vs. male ESCs is unclear. The X-linked ZFX and Y-linked ZFY zinc finger transcription factors regulate autosomal sex-biased gene expression in human female and male fibroblasts and lymphoblastoid cell lines^40^. ZFX has a greater impact than ZFY in regulating sex-biased gene expression in the two cell types^40^. In *in vitro* assays in HeLa cells, the RNA helicases X-linked DDX3X and Y-linked DDX3Y regulate translation of distinct mRNAs due to intrinsic differences in liquid-liquid phase separation propensity^25^. These findings highlight the potential of several conserved X-Y gene pairs to drive sex differences in mammals.

In this study, we show that the *Kdm5c* and *Kdm5d* X-Y gene pair function in a sex hormone independent manner as critical regulators of sex-biased gene expression in mouse ESCs. Previous work has shown that KDM5C regulates different sets of genes in female vs. male mouse brains^32^. KDM5C also negatively regulates bone mass in female but not male mice, which may help explain the increased vulnerability to osteoporosis in females^34^. Additionally, KDM5C regulates adiposity in female mice in a hormone-independent manner^33^. *Kdm5d* was found to regulate sex-biased gene expression in male mouse embryonic fibroblast cells^35^. In these studies, however, the extent to which KDM5C and KDM5D can substitute for each other was not tested. One study did test the requirement of KDM5C and KDM5D and found that the two proteins are redundant in myocardium compaction^41^, suggesting similar functions of the two proteins in the two sexes, although their influence on gene expression was not tested. Our findings suggest that KDM5C and KDM5D function distinctly in the two sexes to regulate sex-biased gene expression in a hormone-independent manner.

Upon the loss of *Kdm5c*, female-biased genes are upregulated in male cells but are downregulated in female cells. The Y chromosome in male cells is one plausible contributor to this opposing effects. Our data suggest that the Y chromosome is necessary to repress a subset of female-biased genes in male ESCs. KDM5C upregulates most expressed single-copy Y-linked genes in male ESCs, including *Ddx3y*, *Eif2s3y*, *Kdm5d*, *Usp9y*, and *Uty*, which may in turn repress female-biased gene expression. Therefore, the reduced expression of these Y-linked repressors of female-biased gene expression may underlie the de-repression of female-biased genes in male ESCs. By contrast, in female cells KDM5C promotes the expression of a subset of female-biased genes. The second X chromosome in female cells is also a contributor to the upregulation of female-biased genes. Our data show that a subset of female-biased genes is upregulated due to the presence of the second active X chromosome in female ESCs. Independently of the second X chromosome, KDM5C upregulates a subset of female-biased genes in female ESCs.

Our results indicate that KDM5C and KDM5D function in overlapping as well as distinct manners to regulate sex-biased gene expression. KDM5C and KDM5D regulate the expression of a subset of sex-biased genes in the same direction. However, *Kdm5c* loss dysregulates a greater number of sex-biased genes relative to *Kdm5d* loss in XY male ESCs. This differential effect may be due in part to the differential levels of *Kdm5c* and *Kdm5d* expression in male ESCs. In XY male ESCs the average expression of *Kdm5c* and *Kdm5c* is 41.6 and 4.6 TPM, respectively. *Kdm5c* and *Kdm5d* expression are also linked, since *Kdm5c* loss reduces *Kdm5d* expression. Despite the lower expression level of *Kdm5d* relative to *Kdm5c*, our data demonstrate a critical regulatory role of KDM5D in WT XY male ESCs. A set of genes are differentially expressed in the absence of *Kdm5d* alone, which suggests that *Kdm5c* cannot compensate for the absence of *Kdm5d*. Moreover, ectopic expression of *Kdm5d* in *X*^5c–fl^*O* female ESCs with intact *Kdm5c* is sufficient to drive a sex-biased gene expression pattern similar to that of WT XY male ESCs. Taken together, these results suggest that KDM5C and KDM5D have diverged in biochemical function. In agreement with this notion, KDM5C contains a larger intrinsically disordered region (IDR) in its C-terminus relative to KDM5D, and only the C-terminal segments are substantially different between the two proteins. Since IDRs can modulate protein-protein interactions^42^, partnering with different proteins could result in the two proteins diverging in their regulatory functions. For example, some genes are targeted by KDM5C but not KDM5D, and vice versa.

KDM5C and KDM5D are thought to function primarily as transcriptional repressors by demethylating the active transcription-associated chromatin marks H3K4me2/3. Moreover, both KDM5C and KDM5D have been shown to interact with the Polycomb Repressive Complex 1.6 (PRC1.6), which deposits the chromatin mark H2AK119ub1 associated with transcriptional repression^30,43^. Consistent with these repressive functions and interactions, we observed greater numbers of upregulated genes compared to downregulated genes in *X*^5c–Δ^*Y*, *XY*^5d–Δ^, and *X*^5c–Δ^*Y*^5d–Δ^ male ESCs relative to WT male ESCs. However, KDM5C has also been shown to function as a transcriptional activator possibly by converting H3K4me2/3 to H3K4me1, which is a signature of transcriptional enhancers^28,36^. Given that KDM5D also demethylates H3K4me2/3 but not H3K4me1^29,43^, it is plausible that KDM5D shares this transcription-activating function. The subset of downregulated sex-biased genes in our mutant cells may reflect the loss of the transcriptional activating functions of KDM5C and KDM5D. Dysregulated sex-biased gene expression may also be due to indirect consequences of *Kdm5c/d* loss.

Unlike many Y-linked genes involved in spermatogenesis, *Kdm5d* is dispensable for male fertility (this study and ref. 44). Rather, our data suggest that a primary role of KDM5D, alongside KDM5C, is shaping sex differences of non-reproductive tissues. Our study underscores the need for further comparative analyses of X-Y gene pairs in non-reproductive tissues to understand their contributions to sex differences in normal physiology and disease susceptibility.

## Acknowledgements

We thank Clair Harris for contributing to the whole genome sequencing mapping. We acknowledge Thomas L. Saunders, Zachary T. Freeman, Elizabeth Hughes, Wanda Filipiak, Galina Gavrilina, and the Transgenic Animal Model Core of the University of Michigan’s Biomedical Research Core Facilities for design and production of *Kdm5d* conditional transgenic mice. Research reported in this publication was supported by the University of Michigan Transgenic Animal Model Core and the Biomedical Research Core Facilities. We acknowledge the services of the University of Michigan Sequencing Core Facility. This work was funded by NIH Institutional Research Service Awards T32-GM007544 (University of Michigan Predoctoral Genetics Training Program; to R.M.M.); T32-HD079342 (University of Michigan Predoctoral Career Training in Reproductive Sciences Program; to R.M.M.); NIH Director’s New Innovator Award (DP2-OD-008646; to S.K.); NIH NICHD R01 Award (R01HD095463; to S.K.); NIH NIGMS R01 Award (R01GM124571; to S.K.); NIH Office of the Director/NICHD TR01 Award (R01HD118514; to S.K.); NIH Awards (R01NS116008 and R01MH133632; to S.I.); and a Farrehi Family Foundation Grant (to S.I.).

## Author contributions

R.M.M., S.K., and S.I. conceived the study and designed the experiments. R.M.M. and M.K.S. derived the ESC lines. R.M.M. characterized, transduced, and analyzed the ESC lines. R.M.M. generated all RNA-seq libraries and performed all analyses. M.K.S. performed FISH experiments. S.I. cloned the *Kdm5d* PHAGE expression vector. R.M.M., S.K., and S.I. wrote the manuscript. All authors have read and agreed to the published version of the manuscript.

## Supplemental information

Document S1. Figures S1–S6

Table S1. Sex hormone nuclear receptor gene expression in all ESC lines

Table S2. List of sgRNAs, single-stranded DNA donor, and primers

## Supplemental figure legends

**Figure S1: Chromosome copy number and pluripotency gene expression in WT *X***^5c–fl^***X***^5c–fl^ **female, WT *X***^5c–fl^***Y* male, and XO ESCs.** (A-C) Chromosome copy number ratios from whole genome sequencing (WGS) of WT *X*^5c–fl^*X*^5c–fl^ female (A), WT *X*^5c–fl^*Y* male (B), and *X*^5c–fl^*O* (C) ESCs. Log_2_ copy number ratios are plotted. Copy number ratios were calculated by dividing the read count in each bin by the mean read count per 1 Mb genome wide. (D) Expression (TPM) of pluripotency genes *Pou5f1*, *Sox2*, *Nanog*, *Zfp42*, *Otx2*, *Pou3f1*, and *T* in WT *X*^5c–fl^*X*^5c–fl^ female, *X*^5c–fl^*O* female, and WT *X*^5c–fl^*Y* male ESCs (mean ± SEM; 3 biological replicates each).

**Figure S2: Generation of *Kdm5d*-mutant mice.** (A) Targeting strategy to generate the *Kdm5d* floxed allele. Mice were bred with germline *Cre* recombinase-containing mice to generate the deleted *Kdm5d* allele (Δ). FP, forward primer; RP, reverse primer. (B) PCR analysis of genomic DNA from male mice harboring the *Kdm5d* WT, floxed, or Δ allele using primers indicated in (A). (C) Genome browser tracks of RNA-seq reads from floxed and *XY*^5d–Δ^ hemizygous mutant male ESCs. Red box, *Kdm5d* exons 11 and 12, which are deleted in *XY*^5d–Δ^ mutant male ESCs.

**Figure S3: Chromosome copy number and pluripotency gene expression in *Kdm5c-* and *Kdm5d*-mutant ESC lines.** (A) Chromosome copy number ratios from whole genome sequencing (WGS) of *X*^5c–fl^*X*^5c–Δ^ heterozygous mutant female, *X*^5c–Δ^*X*^5c–Δ^ homozygous mutant female, *X*^5c–Δ^*Y* hemizygous mutant male, *XY*^5d–Δ^ hemizygous mutant male, and *X*^5c–Δ^ *Y*^5d–Δ^ double-mutant male ESCs. Log_2_ copy number ratios are plotted. Copy number ratios were calculated by dividing the read count in each bin by the mean read count per 1 Mb genome wide. (B) Expression (TPM) of pluripotency genes *Pou5f1*, *Sox2*, *Nanog*, *Zfp42*, *Otx2*, *Pou3f1*, and *T* in WT and mutant female and male ESCs (mean ± SEM; 3 biological replicates each).

**Figure S4: Comparing the relative regulatory contributions of KDM5C and KDM5D.** (A) Overlap of all differentially expressed genes in *X*^5c–Δ^*Y*, *XY*^5d–Δ^, and *X*^5c–Δ^ *Y*^5d–Δ^ mutant male ESCs. (B-G) Genome browser tracks of RNA-seq reads for female-biased genes, *Tspo2* (B), *Xlr3a* (D), and *Egln3* (F), and male-biased genes *Slc44a2* (C), *Creb1* (E), and *Tcerg1l* (G) in WT *X*^5c–fl^*X*^5c–fl^ female and WT *X*^5c–fl^*Y* male ESCs vs. *X*^5c–Δ^*Y* and *XY*^5d–Δ^ mutant male ESCs.

**Figure S5: X- and Y-linked gene expression in *Kdm5c*- and *Kdm5d*-mutant male ESCs.** (A) Heatmap of X-linked gene expression (TPM) in WT *X*^5c–fl^*X*^5c–fl^ female and WT *X*^5c–fl^*Y* male ESCs vs. *X*^5c–Δ^*Y*, *XY*^5d–Δ^, and *X*^5c–Δ^ *Y*^5d-Δ^ mutant male ESCs. Heatmap is clustered by row, scaled by row, and colored by Z-score. (B) Overlap of upregulated X-linked genes in *X*^5c-Δ^*Y* hemizygous mutant male, *XY*^5d-Δ^ hemizygous mutant male, and *X*^5c-Δ^ *Y*^5d-Δ^ double-mutant male ESCs. (C) Heatmap of Y-linked gene expression (TPM) in WT *X*^5c-fl^*Y* male ESCs vs. *X*^5c-Δ^*Y*, *XY*^5d-Δ^, and *X*^5c-Δ^ *Y*^5d-Δ^ mutant male ESCs. Heatmap is clustered by row, scaled by row, and colored by Z-score.

**Figure S6: XX vs. XO quantification in WT *X*^5c-fl^*X*^5c-fl^ and *Kdm5c*-mutant female ESCs.** (A) Table of RNA FISH and WGS data used for X chromosome quantification in all WT *X*^5c-fl^*X*^5c-fl^ and *Kdm5c*-mutant female ESCs. (B) Representative RNA FISH images of an XX nucleus and an XO nucleus. The X chromosome is identified by Xist/Tsix RNA expression (green) and Atrx RNA expression (red). Nuclear DNA is stained with DAPI (blue). Scale bars, 1 µm. (C-D) Formulas used to calculate % X chromosome content by RNA FISH (C) and WGS (D). X, X chromosome. A, autosomes. (E) Correlation of % X chromosome content calculated by RNA FISH vs. WGS. r_s_, Spearman correlation.

## Star methods

### Contact for reagent and resource sharing

Further information and requests for resources and reagents should be directed to and will be fulfilled by the lead contact, Sundeep Kalantry.

### Experimental model and subject details

#### Ethics statement

This study was performed in strict accordance with the recommendations in the Guide for the Care and Use of Laboratory Animals of the National Institutes of Health. All animals were handled according to the protocols approved by the University Committee on Use and Care of Animals (UCUCA) at the University of Michigan (protocols #PRO00010233 and #PRO00011932).

#### Mice

Generation of the *Kdm5c* conditional allele mouse strain has been described previously^37^. The *Kdm5d* (<u>ENSMUSG00000056673</u>) conditional allele mouse strain was generated by the University of Michigan Transgenic Animal Model Core using CRISPR/Cas9 technology. The CRISPOR algorithm^45^ was used to identify a two guide RNA targets (sgRNA) predicted to cut in intronic regions surrounding exons 11 and 12 of *Kdm5d*. sgRNA C201A targets the 5’ region of the exon with 5’ CAAAGCAACTACAAATTATG PAM (GGG) 3’ and C201Y targets the 3’ region with 3’ TAAGAGAACTACATATGTGA PAM (TGG) 5’. Phosphorothioate modified sgRNAs were synthesized by Synthego. The sgRNAs (30 ng/ul) were complexed with enhanced specificity Cas9 proteins (eSpCas9, 50 ng/ul; Millipore-Sigma, #ESPCAS9PRO)^46^ and individually tested to determine if the ribonucleoprotein (RNP) complexes caused chromosome breaks in mouse zygotes. RNPs were microinjected into zygotes, which were placed in culture until they developed into blastocysts. DNA was extracted from individual blastocysts for analysis. PCR with primers spanning the predicted cut site was used to generate amplicons for Sanger sequencing^47^. Amplicons were produced for both C201A with forward primer: 5’ AACAAATGGGATAAAATAGCTTAACACTAGCA 3’ and reverse primer 5’: GCATGCATTCCTGTACATAATAAGTGTCT, 3’; 520 bp amplicon, and C201Y with forward primer: 5’ TTATTCCATTAACTACCTGCACTGGTGA 3’ and reverse primer 5’: ACCAGAAAATATCAGAACAAGAGTGAGC 3’; 772 bp amplicon. Sequencing electropherograms of amplicons from individual blastocysts were evaluated to determine if small insertions/deletions caused by non-homologous end joining (NHEJ) repair of chromosome breaks were present^48^. sgRNA C201A and C201Y induced chromosome breaks and had high activity scores^49^. Active RNPs were mixed with a synthetic long single-stranded DNA donor (ssDNA donor, 5 ng/ul; IDT; Table S2) prior to microinjection into fertilized mouse zygotes produced by mating superovulated C57BL/6:SJL (B6SJLF1) F1 hybrid female mice (Jackson Laboratory stock no. 100012) with B6SJLF1 male mice as described^50^, and gave rise to founder mice. Both the *Kdm5c* and *Kdm5d* conditional allele strains were maintained on mixed backgrounds. To genotype the mice, ear punches were lysed in tail lysis buffer (50 mM KCL, 10 mM Tris-Cl (pH 8.3), 2.5 mM MgCl_2_, 0.1 mg/mL gelatin, 0.45% NP-40, and 0.45% Tween-20). The ear punch and lysis buffer mixture were incubated at 56°C overnight with Proteinase K (Invitrogen, #4333793). The following day the samples were cooked at ≥85°C for 10 minutes to inactivate the Proteinase K. PCR reactions were carried out using standard conditions. The following primers were used to genotype the *Kdm5d* ^fl^ and *Kdm5d* ^Δ^ mice: forward primer #1 5’ TTAGAAAGAGTTCTTACTGGGT 3’, forward primer #2 5’ TCGCTAGCAGCAGAACACTT 3’, reverse primer #1 5’ AATGTGATGCCAGTGCTGGA 3’.

#### ESC derivation

Embryonic stem cells (ESCs) were derived by following a published protocol^51^. Briefly, pregnant females were identified by copulatory plug checking and were dissected 3.5 days after plugging. Pregnant females were euthanized by cervical dislocation, and uterine horns were dissected out using sterile protocol. Uterine horns were flushed using 3 mL syringes (Thermo Fisher Scientific, #22-257-120) equipped with 25-gauge needles (Thermo Fisher Scientific, #14-826AA) filled with sterile media to remove pre-implantation blastocysts. Blastocysts were transferred into prepared 4-well tissue culture dishes (Thermo Fisher Scientific, #1256572) containing mouse embryonic fibroblast feeder cells and ESC derivation media. Mouse embryonic fibroblasts (MEFs) were derived from E14.5 embryos and were mitotically inactivated by irradiation. ESC derivation media is composed of KnockOut DMEM (Invitrogen, #10829018) base with 20% KnockOut serum replacement (Invitrogen, #10828028), 1x Penicillin-streptomycin (Invitrogen, #15140122), 2 mM L-Glutamine (Invitrogen, #25030081), 1x non-essential amino acids (Invitrogen, #11140050), and 0.1 mM 2-mercaptoethanol (Millipore-Sigma, #M3148). Media was supplemented with 3 µM CHIR99021 (Millipore-Sigma, # SML1046), 1µM PD0325901 (Millipore-Sigma, #PZ0162), and 10^3^ units/mL of LIF (R&D Systems, #8878-LF-100/CF). Embryos were incubated at 37°C at 5% CO_2_, with media being changed every other day until outgrowths were large and ready to be dissociated into single cells. Around day 4-6 in culture, outgrowths were dissociated into single cells in 0.05% trypsin-EDTA (Invitrogen, #25300062) and transferred into 96-well wells of MEFs. Culture was then progressively scaled up with media being changed daily, and passaging occurring every 2-3 days.

#### ESC culture

ESCs were maintained in culture following a published protocol^51^. Briefly, ESCs were maintained on MEFs in media with a KnockOut DMEM base (Invitrogen, #10829018) containing 15% KnockOut Serum replacement (Invitrogen, #10828028), 5% embryonic stem cell qualified fetal bovine serum (R&D Systems, #S10250), 1x Penicillin-streptomycin (Invitrogen, #15140122), 2 mM L-Glutamine (Invitrogen, #25030081), 1x non-essential amino acids (Invitrogen, #11140050), and 0.1 mM 2-mercaptoethanol (Millipore-Sigma, #M3148). Media was supplemented with 10^3^ units/mL of LIF (R&D Systems, #8878-LF-100/CF). Cells were cultured at 37°C at 5% CO_2_ with media changed daily and were passaged every 2-3 days using 0.05% trypsin-EDTA (Invitrogen, #25300062).

### Generating *X*^5c-fl^*O* female ESC lines

To generate *X*^5c-fl^*O* female ESC lines, we subcloned XO female ESCs from an *X*^5c-fl^*X*^5c-Δ^ parental ESC line; upon prolonged culture XX ESC lines tend to lose an X chromosome^38,52^. XO status was verified by RNA fluorescence in situ hybridization (FISH; Methods) for expressed X-linked genes (Figure S5).

#### Lentiviral transduction in ESCs

To generate plasmids for constitutive expression, mouse *Kdm5d* cDNA was cloned into an in-house generated PGK promoter-driven Strep-tag II lentiviral vector by Gateway LR recombination. To produce the lentivirus, HEK293T cells were grown to 50-70% confluency in a 10 cm dish and co-transfected with 6 µg of the lentiviral construct, 3 µg of the psPAX2 packaging plasmid (Addgene plasmid, #12260), and 1.5 µg of the pMD2.G envelope plasmid (Addgene plasmid, #12259) with Lipofectamine 2000 (Invitrogen, #11668). Lentiviral particles were collected at 48- and 72-hours post-transfection and concentrated using Lenti-X Concentrator (Takara Bio, #631231). ESCs were transduced with the concentrated lentivirus and 10 µg/mL Polybrene (Millipore-Sigma, #TR-1003-G) and selected with 2 µg/mL Puromycin (Millipore-Sigma, #P8833-25MG) beginning 24 hours after transduction. Resistant ESC colonies were sub-cloned to generate clonal cell lines.

## Method details

### Genomic DNA isolation and low-pass whole genome sequencing

Genomic DNA was isolated from cell pellets using the Monarch Genomic DNA Purification Kit (NEB, #T3010L) following the manufacturer’s protocol for cultured cells. Libraries were prepared by the University of Michigan Advanced Genomics Core (AGC) using the NEBNext Ultra II FS DNA Library Prep Kit (NEB, #E7805) following the manufacturer’s instructions. Pooled libraries were sequenced using 150 base pair paired-end sequencing on the Illumina NovaSeq X Plus by the University of Michigan AGC.

### RNA isolation and RNA-seq library preparation

Total RNA was isolated from cell pellets by TRIzol (Invitrogen, #15596-018) extraction following the manufacturer’s protocol. RNA concentration was assessed using the Qubit RNA HS assay (Invitrogen, #Q32852) with a Qubit 4.0 fluorometer (Thermo Fisher Scientific). RNA quality (RIN^e^) was assessed with Agilent High Sensitivity RNA ScreenTape analysis (Agilent Technologies; ScreenTape, #5067-5579; ScreenTape Sample Buffer, #5067-5580; ScreenTape Ladder, #5067-5581) using the TapeStation Analysis Software 3.2 on a TapeStation 4150 instrument (Agilent Technologies). Prior to mRNA purification, 2 µL of a 1:100 dilution of ERCC ExFold RNA Spike-In Mix 1 or Mix 2 (Thermo Fisher Scientific, #44-567-39) were added to 1 µg of total RNA. mRNA purification was performed using the NEBNext Poly(A) mRNA Magnetic Isolation Module (NEB, #E7490L) following the manufacturer’s instructions. Libraries were prepared using purified mRNA using the NEBNext Ultra II Directional RNA Library Prep Kit for Illumina (NEB, #E7765L) following the manufacturer’s instructions. Pooled libraries were sequenced using 150 base pair paired-end sequencing on the NovaSeq 6000 at the University of Michigan AGC.

### RNA fluorescence *in situ* hybridization (FISH)

RNA FISH with double-stranded (ds) probes was performed following a published protocol^53^. Briefly, dsRNA FISH probes for Xist/Tsix RNA and Atrx RNA were labeled separately using the BioPrime DNA Labeling System (Invitrogen, #18094011). Probes were randomly primed and extended by Klenow fragment in the presence of Cy3-dCTP (Atrx RNA; Millipore-Sigma, #GEPA53021) or Fluorescein 12 (FITC)-dUTP (Xist RNA; Thermo Fisher Scientific, #FERR0101). After labeling, probes were purified using ProbeQuant G-50 Micro Columns (Cytiva, #28903408) and precipitated with 0.3M sodium acetate (Thermo Fisher Scientific, #50-843-082) and 300 µg of yeast tRNA (Invitrogen, #15401011) in 100% molecular biology grade ethanol (Thermo Fisher Scientific, #BP2818-500). Xist/Tsix RNA and Atrx RNA probes were then precipitated together with 300 µg yeast tRNA, 0.3M sodium acetate, and 150 µg of sheared boiled salmon sperm (Invitrogen, #15632-011). Probes were pelleted by centrifuging at 21,130 xg for 20 minutes at 4°C and subsequently washed consecutively with 70% ethanol and 100% ethanol. Pellets were dried and resuspended in 100% deionized formamide (Thermo Fisher Scientific, #50-256-002). Probes were denatured in formamide by incubating at 90°C for 10 minutes, followed by 5 minutes on ice. A hybridization solution consisting of 4x SSC (Invitrogen, #AM9765), 20% dextran sulfate (Millipore-Sigma, #3730100ML), and water was then added to the denatured probes.

For the quantification of XX vs. XO nuclei in female ESC lines, Xist/Tsix RNA and Atrx RNA FISH were performed concurrently using dsRNA FISH probes. ESCs were grown on gelatinized coverslips and permeabilized through sequential treatment with ice-cold cytoskeletal extraction buffer (CSK:100 mM NaCl, 300 mM sucrose, 3 mM MgCl2, 10 mM PIPES buffer, pH 6.8) without followed by with 0.4% Triton X-100 (Thermo Fisher Scientific, #BP151-100). Cells incubated in 4% paraformaldehyde (Electron Microscopy Sciences, #15710) for 10 minutes to fix to the coverslips. Fixed cells were then dehydrated through 2-minute sequential incubations in 70%, 85%, 95%, and 100% ethanol and subsequently air dried. Coverslips were hybridized to the probe overnight at 37°C in a humid chamber. Coverslips were washed sequentially in a pre-warmed 2x SSC/50% formamide solution, a 2x SSC solution, and a 1x SSC solution at 39°C, with three 7-minute washes per solution. DAPI (Invitrogen, #D21490) was added during the third 2x SSC wash. Coverslips were mounted on slides with Vectashield (Vector Labs, #H-1000-10).

### RT-qPCR

Total RNA was isolated from cell pellets by TRIzol (Invitrogen, #15596-018) extraction following the manufacturer’s protocol. RT-qPCR reactions were carried out using the Luna Universal One-Step RT-qPCR Kit (NEB, #E3005L). Quantification was performed using a SYBR Green-based method on an Eppendorf Realplex Mastercycler (software version 2.2.0.84). The housekeeping gene *Tbp* was used as an internal control for normalization. The following primers were used for RT-qPCR: *Tbp*: forward 5’ TTCAGAGGATGCTCTAGGGAAGA 3’, reverse 5’ CTGTGGAGTAAGTCCTGTGCC 3’; *Kdm5d* transgene: forward in Strep-tag II 5’ GTGGAGCCACCCCCAGTTC 3’, reverse in *Kdm5d* exon 1 5’ GAAAGTCGTCAGATCCTGGCTTC 3’; total *Kdm5d*: forward in *Kdm5d* exon 10 5’ GCAGCATTGAGGAGGATGTG 3’, reverse in *Kdm5d* exon 11 5’ CCCAGTGCAGGTAGTTAATGG 3’.

### Quantification and statistical analysis <u>Whole genome sequencing data processing</u>

Raw sequencing fastq data was adaptor-trimmed using the trimmomatic software (v0.36) with the settings ILLUMINACLIP:“+Trim_FA+”:2:30:10:5:true SLIDINGWINDOW:5:30. Trimmed data was aligned to the mouse mm9 (MGSCv37) genome assembly using STAR (v2.5.2a) with the options --outFilterMismatchNmax 10 and --outFilterMultimapNmax 10. SAM files were converted to BAM files using the samtools software (v0.1.19). A Linux shell command labeled each read pair based on genomic position. Reads per 1 Mb bin across the genome were counted by the groupby command using the Bedtools software (v2.30.0). Copy number ratios for each 1 Mb bin were calculated by dividing the number of counts in a bin by the average number of counts per 1 Mb in that sample. The R program ggplot2 (v3.4.2) was used to generate scatterplots of chromosome copy number ratios per 1 Mb bin across genomic coordinates.

Due to reduced mappability of the sex chromosomes^54^, in females, the copy number ratio of the X chromosome (X) is not equivalent to that of autosomes (A; i.e. 1), and in males, the X:A ratio is not exactly 0.5. To account for the reduced mappability of the X, we normalized mean X:A copy number ratios in WT and mutant female ESCs to mean X:A ratios from WT XX mouse organ WGS sequencing (Mean X:A counts per 1 Mb in WT XX mouse organs = 0.8676; Figure S6A and D).

### RNA-seq data processing and quality control

Quality control analysis was conducted using FastQC (v0.11.9). Raw sequencing fastq data was adaptor-trimmed using the trimmomatic software (v0.36) with the settings ILLUMINACLIP:TruSeq3-PE.fa:2:30:10:5:true SLIDINGWINDOW:5:30. Trimmed data was aligned to the mouse mm9 (MGSCv37) genome assembly using STAR (v2.5.2a) with the options --outFilterMismatchNmax 10 and --outFilterMultimapNmax 1. SAM files were converted to BAM files, sorted by coordinate, and indexed using the samtools software (v0.1.19). ERCC Spike-in sequence annotations were merged with the mm9 transcript annotation file for downstream read count analysis. Reads were aligned to exons defined by RefSeq and counted by the Python package HTseq by running the command HTSeq.scripts.count (Python v3.8). Read counts were used to calculate transcripts per million (TPM) to quantify gene expression level. The R package erccdashboard (v1.38.0) was used to estimate the dynamic range and diagnostic performance of the RNA-seq.

### RNA-seq analysis and data visualization

Gene counts were normalized and used to identify differentially expressed genes using edgeR (v3.13) in R (v4.2.3) with the cutoffs of Log_2_FC > 1, FDR < 0.01, and CPM ≥ 1. *P* values were corrected for multiple hypothesis testing by the Benjamini-Hochberg method. The program ggplot2 (v3.4.2) was used to generate MA plots, scatterplots, bar plots, boxplots, and histograms. Heatmaps were generated using pheatmap (v1.0.12). Venn diagrams were generated using VennDiagram (v1.7.3). Mouse KDM5C UniProt accession #P41230 and mouse KDM5D UniProt accession #Q62240 were referenced to generate Figure 2A-B. Genome browser RNA-sequencing profiles were generated using the combination of packages: karyoploteR (v1.22.0), rtracklayer (v1.56.1), GenomicRanges (v1.48.0), diffloop (1.24.0) and TxDb.Mmusculus.UCSC.mm9.knownGene (v3.2.2).

### Statistical analysis

Binomial logistic regressions, Chi-squared tests, and Spearman correlation analysis were all performed using the R package stats (4.4.1). Binomial logistic regressions were performed to compare the overlap between differentially expressed genes and sex-biased genes. Chi-squared tests were performed to determine the expected numbers of female- and male-biased genes per chromosome in WT ESCs (Figure 1C and D). Spearman correlation analysis was used to calculate the spearman rank coefficient, rho (r_s_), for Figure 3I and Figure S3E. Linear models were generated using the R package nlme (3.1-162) for Figure 3I. Kruskal-wallis tests were performed on the data from Figure S1D and Figure S3B using the online program (https://www.statskingdom.com/kruskal-wallis-calculator.html) with Bonferroni correction.

## Data Availability

The RNA-seq data are available in NCBI’s Gene Expression Omnibus (Accession number: GSE277494). The WGS raw data are deposited to SRA under the BioProject Accession number: PRJNA1162514.

## Key resources table

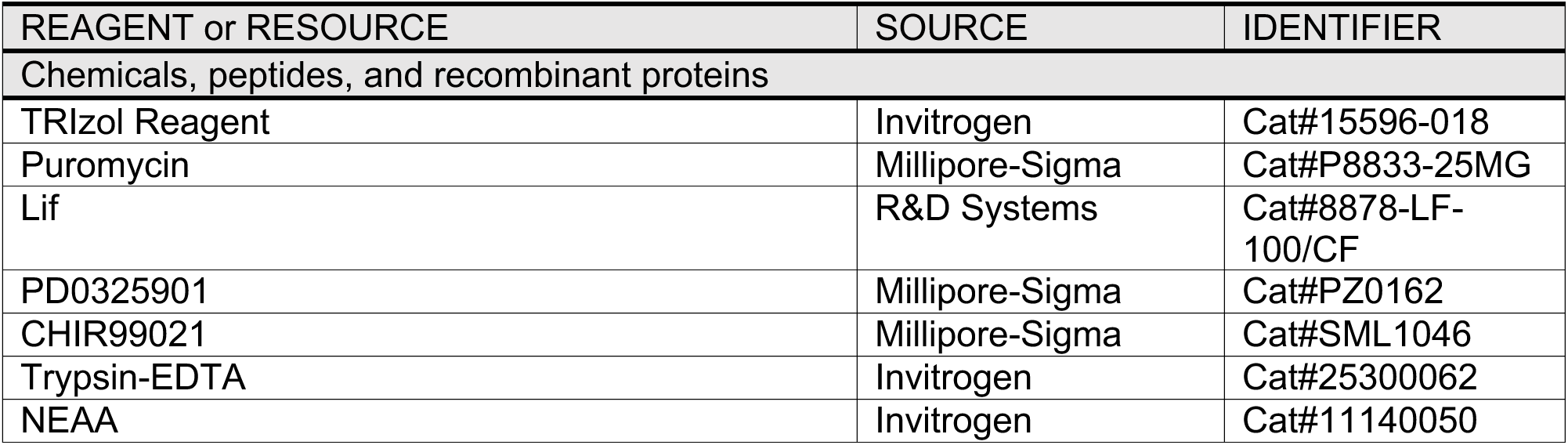
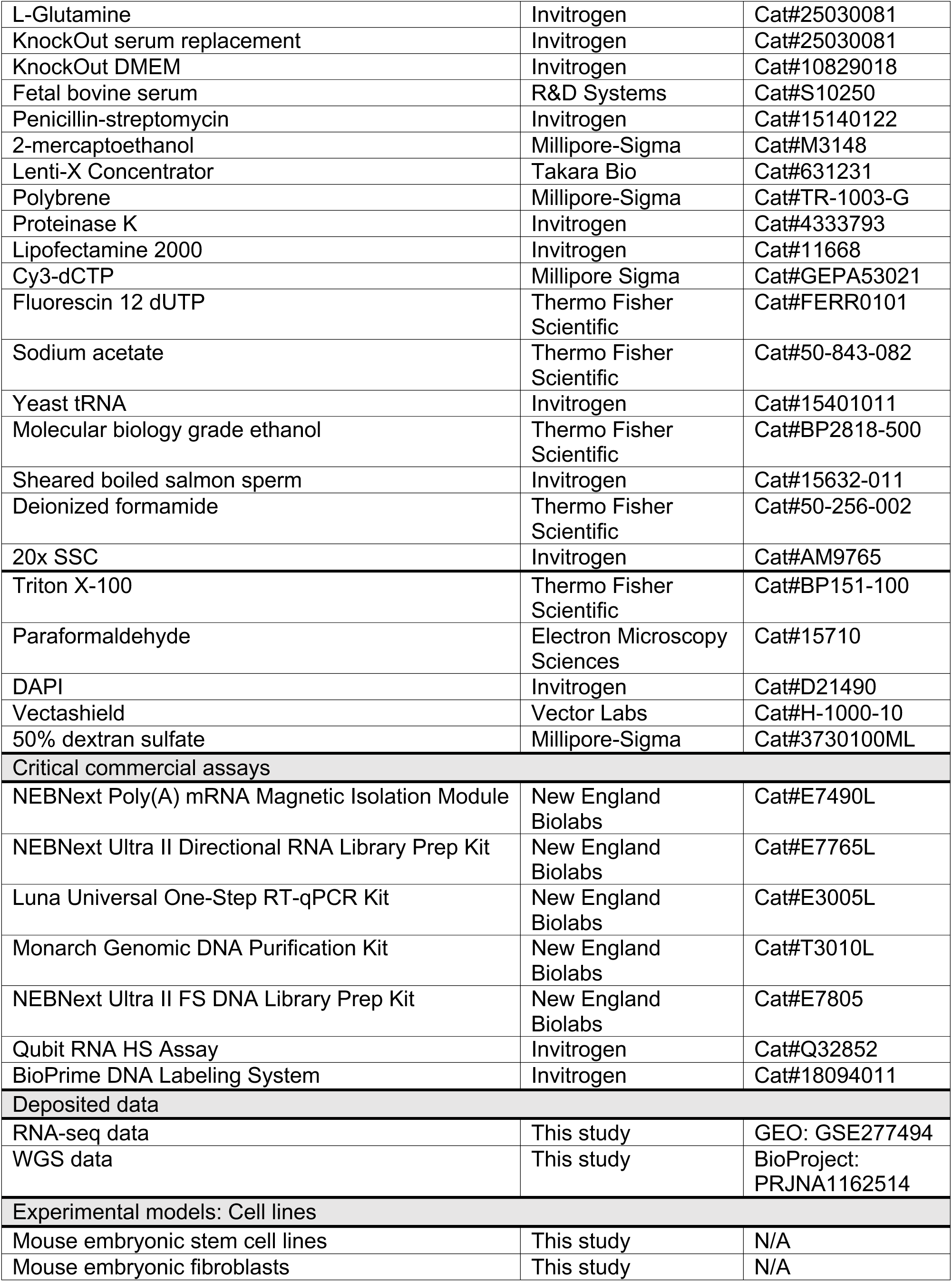
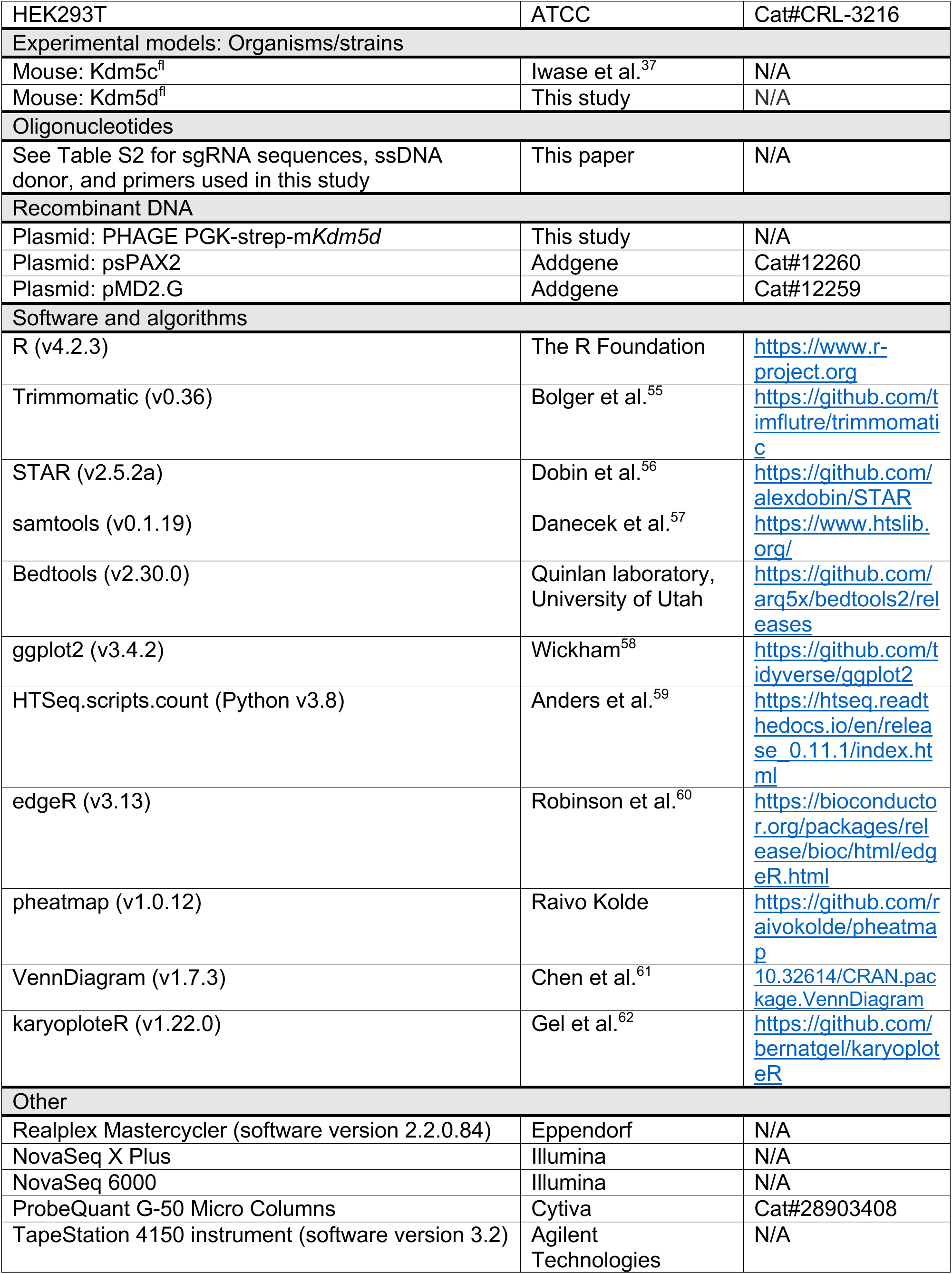
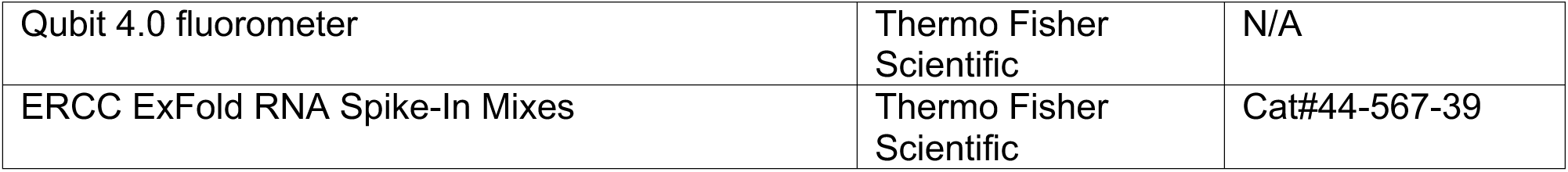

