## Supplementary material for "Regulation of Sex-biased Gene Expression by the Ancestral X-Y Chromosomal Gene Pair *Kdm5c-Kdm5d*": Document S1. Figures S1-S6

### Figure S1

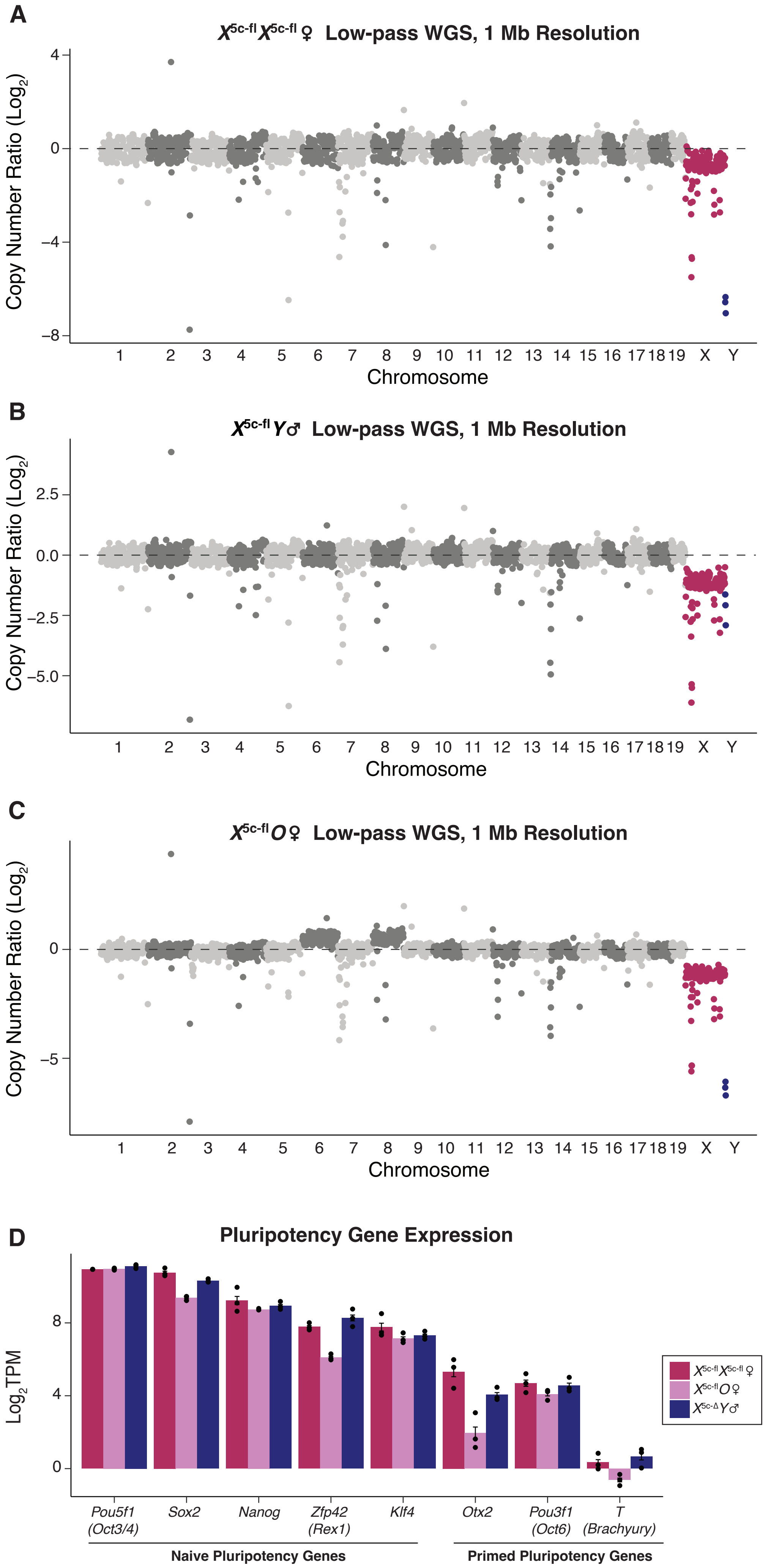

A
$X^{5c\text{-}fl}X^{5c\text{-}fl}$♀ Low-pass WGS, 1 Mb Resolution
Copy Number Ratio (Log2)
4
0
−4
−8
1 2 3 4 5 6 7 8 9 10 11 12 13 14 15 16 17 18 19 X Y
Chromosome
B
$X^{5c\text{-}fl}Y$♂ Low-pass WGS, 1 Mb Resolution
Copy Number Ratio (Log2)
2.5
0.0
−2.5
−5.0
1 2 3 4 5 6 7 8 9 10 11 12 13 14 15 16 17 18 19 X Y
Chromosome
C
$X^{5c\text{-}fl}O$♀ Low-pass WGS, 1 Mb Resolution
Copy Number Ratio (Log2)
0
−5
1 2 3 4 5 6 7 8 9 10 11 12 13 14 15 16 17 18 19 X Y
Chromosome
D
Pluripotency Gene Expression
Log2TPM
8
4
0
Pou5f1 (Oct3/4)
Sox2
Nanog
Zfp42 (Rex1)
Klf4
Otx2
Pou3f1 (Oct6)
T (Brachyury)
Naive Pluripotency Genes
Primed Pluripotency Genes
$X^{5c\text{-}fl}X^{5c\text{-}fl}$♀
$X^{5c\text{-}fl}O$♀
$X^{5c\text{-}\Delta}Y$♂

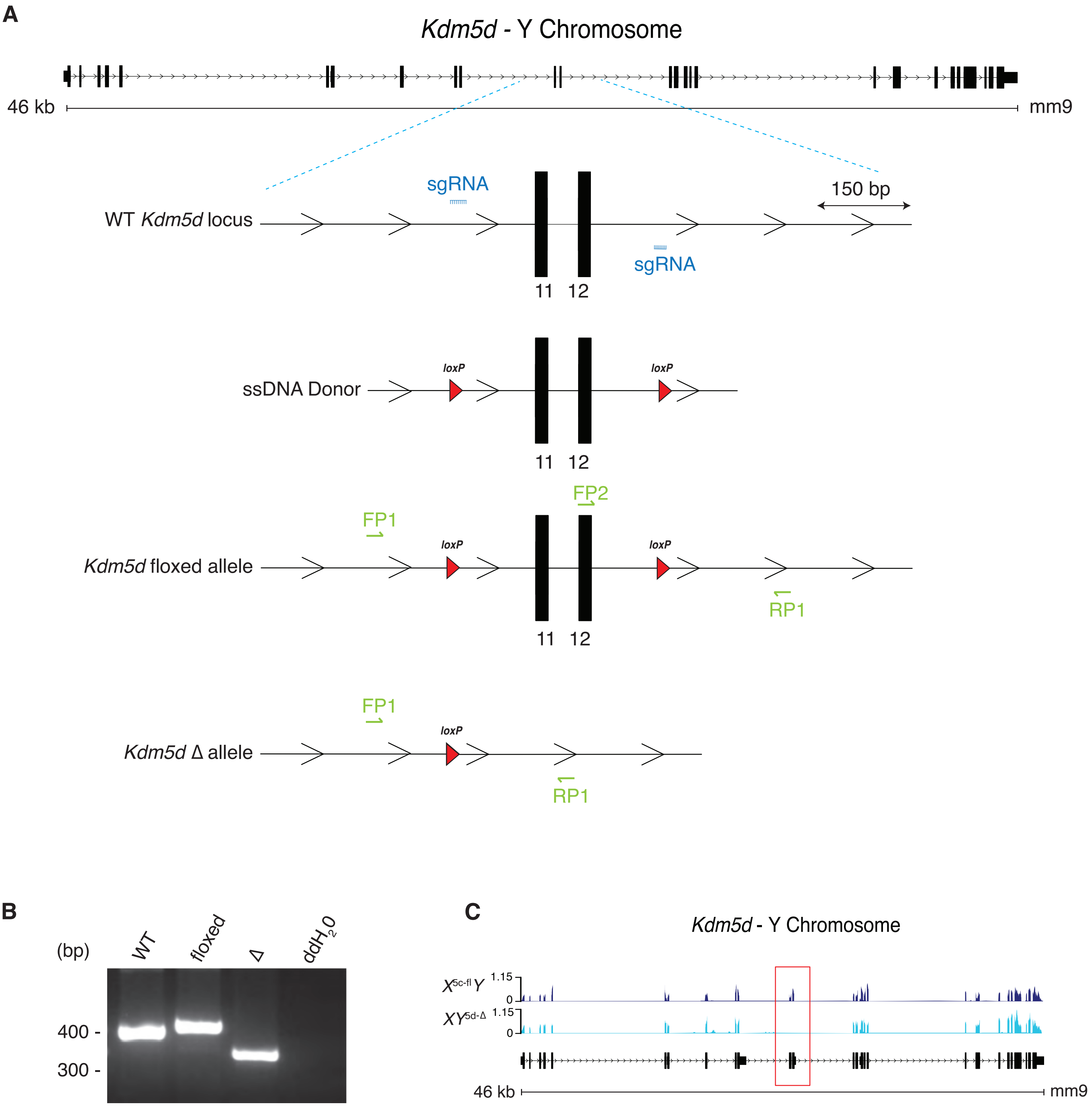
A
Kdm5d - Y Chromosome
46 kb
mm9
sgRNA
150 bp
WT Kdm5d locus
sgRNA
11 12
loxP
ssDNA Donor
loxP
11 12
FP2
FP1
loxP
loxP
Kdm5d floxed allele
RP1
11 12
FP1
loxP
Kdm5d Δ allele
RP1
B
(bp)
WT
floxed
Δ
ddH2O
400 -
300 -
C
Kdm5d - Y Chromosome
X5c-fl Y 1.15
0
XY5d-Δ 1.15
0
46 kb
mm9

**Figure S3**

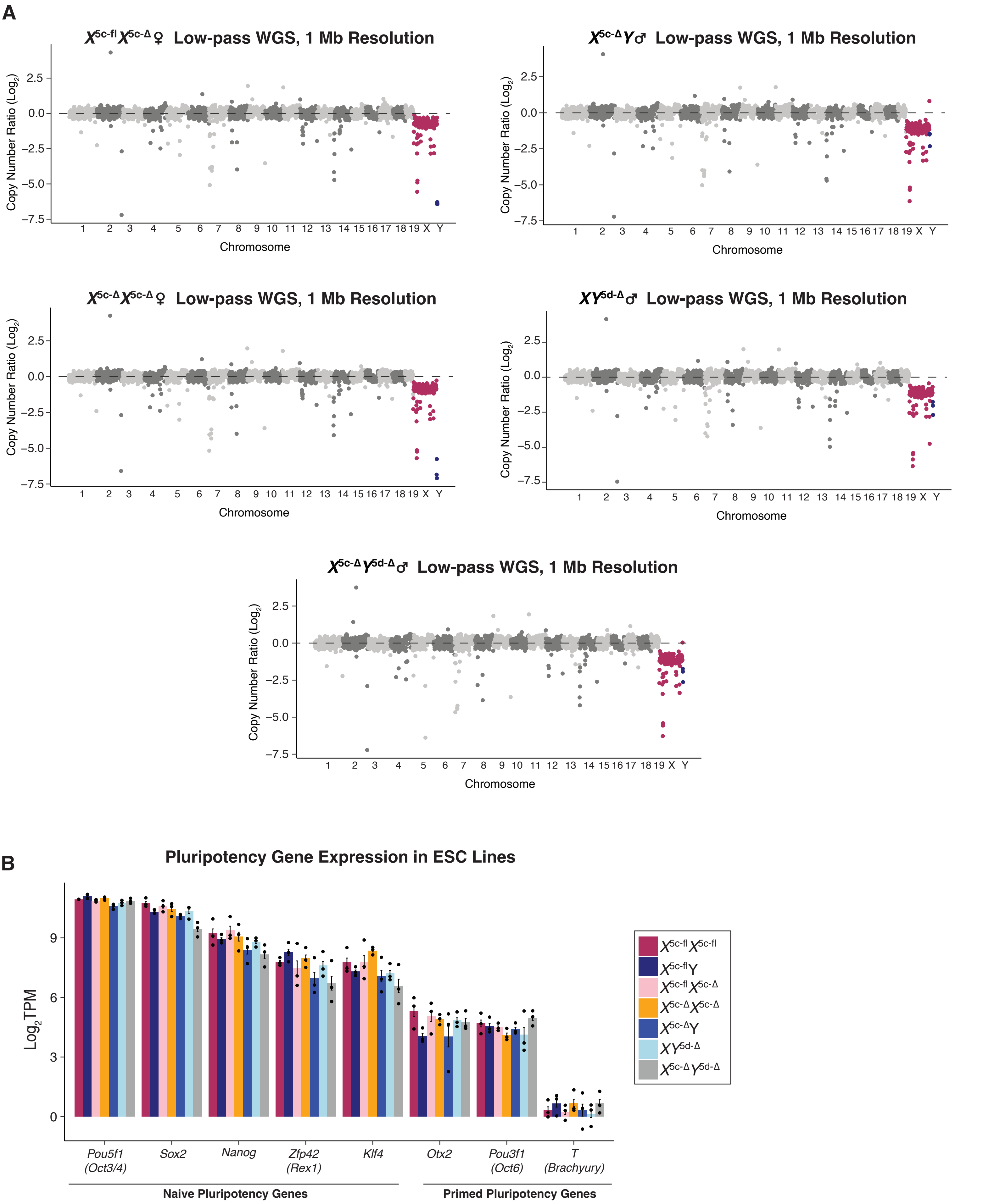
A
X5c-fl X5c-Δ ♀ Low-pass WGS, 1 Mb Resolution
X5c-Δ Y♂ Low-pass WGS, 1 Mb Resolution
X5c-Δ X5c-Δ ♀ Low-pass WGS, 1 Mb Resolution
XY5d-Δ ♂ Low-pass WGS, 1 Mb Resolution
X5c-Δ Y5d-Δ ♂ Low-pass WGS, 1 Mb Resolution
Copy Number Ratio (Log2)
Chromosome
1 2 3 4 5 6 7 8 9 10 11 12 13 14 15 16 17 18 19 X Y
2.5
0.0
−2.5
−5.0
−7.5
B
Pluripotency Gene Expression in ESC Lines
Log2 TPM
0
3
6
9
Pou5f1 (Oct3/4)
Sox2
Nanog
Zfp42 (Rex1)
Klf4
Otx2
Pou3f1 (Oct6)
T (Brachyury)
Naive Pluripotency Genes
Primed Pluripotency Genes
X5c-fl X5c-fl
X5c-fl Y
X5c-fl X5c-Δ
X5c-Δ X5c-Δ
X5c-Δ Y
XY5d-Δ
X5c-Δ Y5d-Δ

**Figure S4**

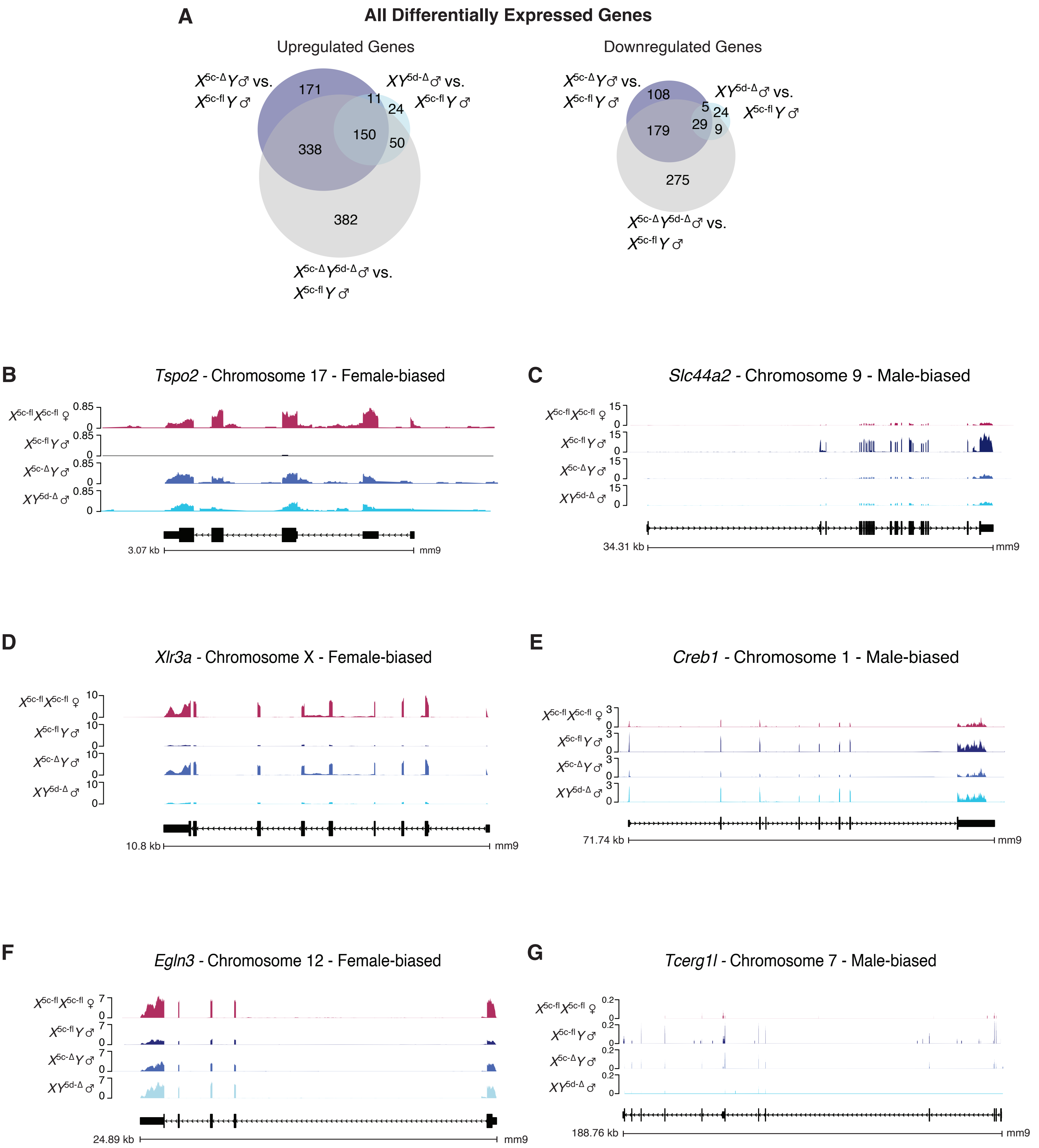
A
All Differentially Expressed Genes
Upregulated Genes
X5c-Δ Y♂ vs. X5c-fl Y♂
XY5d-Δ♂ vs. X5c-fl Y♂
X5c-Δ Y5d-Δ♂ vs. X5c-fl Y♂
171
11
24
150
338
50
382
Downregulated Genes
108
5
24
29
9
179
275
B
Tspo2 - Chromosome 17 - Female-biased
3.07 kb
mm9
C
Slc44a2 - Chromosome 9 - Male-biased
34.31 kb
mm9
D
Xlr3a - Chromosome X - Female-biased
10.8 kb
mm9
E
Creb1 - Chromosome 1 - Male-biased
71.74 kb
mm9
F
Egln3 - Chromosome 12 - Female-biased
24.89 kb
mm9
G
Tcerg1l - Chromosome 7 - Male-biased
188.76 kb
mm9
X5c-fl X5c-fl ♀
X5c-fl Y♂
X5c-Δ Y♂
XY5d-Δ ♂

**Figure S5**

**A** **All Expressed X-linked Genes (n = 466)**

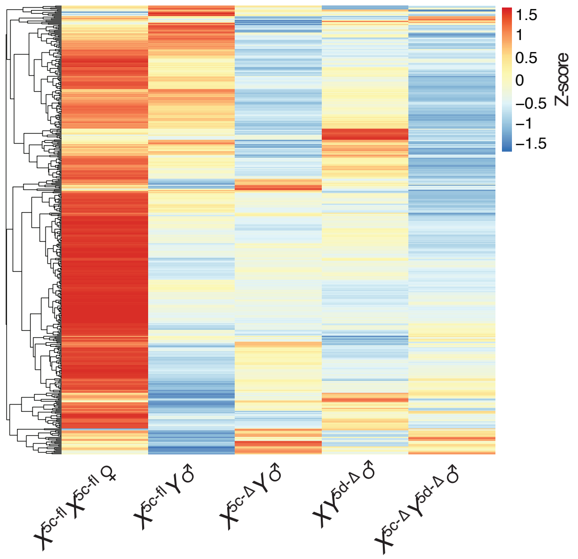


**B** **Upregulated X-linked Genes**

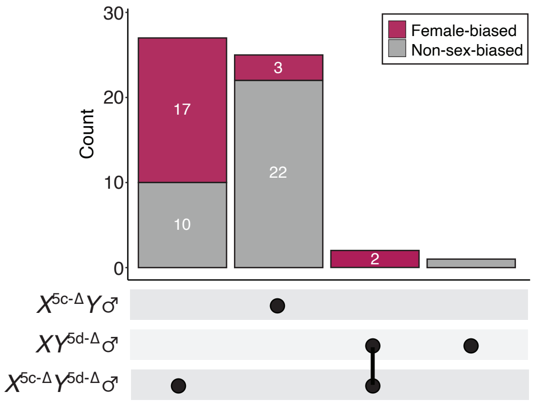


**C** **All Expressed Y-linked Genes**

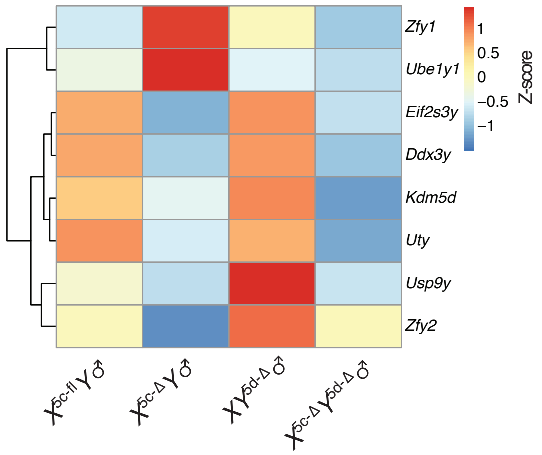

**A** **X Chromosome Quantification in Female ESC Lines**

| Cell line | RNA-FISH Quantification: % XX nuclei | % XO nuclei | # nuclei counted | %X chr content | WGS Quantification: Avg. A counts/1Mb | Avg. X counts/1Mb | %X:A | %X rel.to XX |
|---|---|---|---|---|---|---|---|---|
| $X^{5c\text{-}fl}X^{5c\text{-}fl}$♀ #1 | 70% | 30% | 257 | 85% | 3134.09 | 2198.07 | 70% | 81% |
| $X^{5c\text{-}fl}X^{5c\text{-}fl}$♀ #2 | 84% | 16% | 295 | 92% | 3379.63 | 2248.38 | 67% | 77% |
| $X^{5c\text{-}fl}X^{5c\text{-}fl}$♀ #3 | 70% | 30% | 167 | 85% | 2973.08 | 1754.95 | 59% | 68% |
| $X^{5c\text{-}fl}X^{5c\text{-}\Delta}$♀ #1 | 74% | 26% | 168 | 87% | 4317.76 | 2394.68 | 55% | 64% |
| $X^{5c\text{-}fl}X^{5c\text{-}\Delta}$♀ #2 | 95% | 5% | 260 | 97.5% | 3082.44 | 1847.20 | 60% | 69% |
| $X^{5c\text{-}fl}X^{5c\text{-}\Delta}$♀ #3 | 74% | 26% | 242 | 87% | 2853.83 | 1592.36 | 56% | 64% |
| $X^{5c\text{-}\Delta}X^{5c\text{-}\Delta}$♀ #1 | 20% | 80% | 810 | 60% | 2683.57 | 1349.73 | 50% | 58% |
| $X^{5c\text{-}\Delta}X^{5c\text{-}\Delta}$♀ #2 | 30% | 70% | 490 | 65% | 2708.57 | 1171.29 | 43% | 50% |
| $X^{5c\text{-}\Delta}X^{5c\text{-}\Delta}$♀ #3 | 35% | 65% | 748 | 67.5% | 2666.19 | 1338.70 | 50% | 58% |

**B** **RNA FISH for XX vs. XO**

XX nucleus XO nucleus

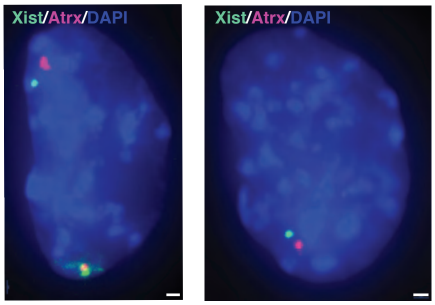


**C** **RNA FISH % X Chromosome Content Formula**

$$\text{RNA FISH \% X chr content} = \frac{2(\%XX) + 1(\%XO)}{200}$$

**D** **WGS % X Chromosome Content Formula**

$$\text{WGS \% X chr content} = \frac{\left(\dfrac{\text{Avg. (X counts/1Mb) in ESC line}}{\text{Avg. (A counts/1Mb) in ESC line}}\right)}{\left(\dfrac{\text{Avg. (X counts/1Mb) in XX mouse}}{\text{Avg. (A counts/1Mb) in XX mouse}}\right)}$$

**E** **Correlation of RNA FISH vs. WGS**

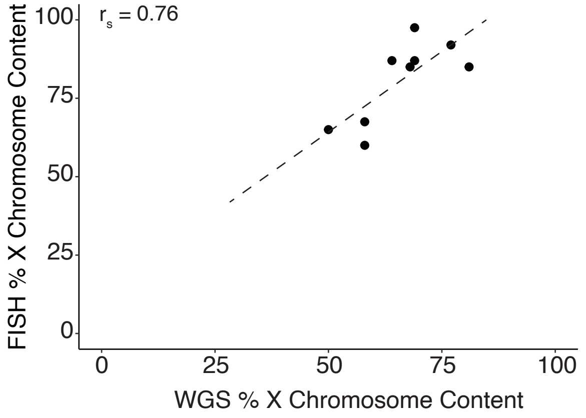
