## Supplementary material for "Regulation of Sex-biased Gene Expression by the Ancestral X-Y Chromosomal Gene Pair *Kdm5c-Kdm5d*": Table S2

**Table S2: Sequences of all oligonucleotides used in this study**

| **sgRNA sequences** | | |
|---|---|---|
| **Name** | **Source** | **Sequence** |
| sgRNA C201A | Synthego | 5’ CAAAGCAACTACAAATTATG 3’ |
| sgRNA C201Y | Synthego | 5’ TCACATATGTAGTTCTCTTA 3’ |

| **Synthetic ssDNA donor sequence[a]** | | |
|---|---|---|
| **Name[b]** | **Source** | **Sequence** |
| *Kdm5d* floxed exons 11 and 12 donor | IDT | 5’cactagcatttagaaagagttcttactgggtcatatgaaccttcatttgcattattacttttaacatgattatacttttgtagaaaagaaaaattacatttacaaatcatacatatacttccaaagcaact**ataacttcgtataatgtatgctatacgaagttat**acaaattatggggtgcattatattctgagaacacagatatttttgtctgtagatctggctcattttacctttcttagtccaatctcactccctacctacaatgtaattctgtgactcaacagGAATATGCTGCTTGTGGTTGGAATCTCAATGTGATGCCAGTGCTGGATCAGTCTGTTCTCTGCCACATCAATGCAGACATCTCAGGCATGAAAGTGCCCTGGTTATATGTGGGCATGGTGTTTTCAGCATTTTGTTGGCATATTGAGGATCACTGGAGTTATTCCATTAACTACCTGCACTGgtgagtgatacaccaataactagaatgtatcaagacacttattatgtacaggaatgcatgcaaactttatatatctcttttctctggcagGGGTGAACCAAAGACCTGGTATGGAGTACCTTCGCTAGCAGCAGAACACTTAGAGGATGTAATGAAGAGACTTACACCAGAGTTGTTTGACAGCCAACCTGACCTCCTGCACCAACTTGTCACTCTGATGAATCCTAACACTCTCATGTCACATGGAGTGCCAgtgagtatctggaaatgaggctcaaggggaaaatgcagctttttcaaaatatgtgcatgtgcacacagaaaggaatgttttttctgactattgacagaccatcacatatgttaag**ataacttcgtataatgtatgctatacgaagttat**ttctcttacagtgcttctagcaaagttatagaaaaaagaaaaaacagaaataggattgttatgtaacagcataaaaagatcttaaaaacctagtctagaaccag 3’ |

| **Primer sequences** | | | |
|---|---|---|---|
| **Gene/region amplified[b]** | **Purpose** | **Forward Primer Sequence** | **Reverse Primer Sequence** |
| C201A cut site | Indel screening | 5’AACAAATGGGATAAAATAGCTTAACACTAGCA 3’ | 5’GCATGCATTCCTGTACATAATAAGTGTCT 3’ |
| C201Y cut site | Indel screening | 5’TTATTCCATTAACTACCTGCACTGGTGA 3’ | 5’ACCAGAAAATATCAGAACAAGAGTGAGC 3’ |
| Kdm5d fl vs. $\Delta$ | mouse genotyping | 5’TTAGAAAGAGTTCTTACTGGGT 3’<br>5’TCGCTAGCAGCAGAACACTT 3’ | 5’AATGTGATGCCAGTGCTGGA 3’ |
| *Tbp* | RT-qPCR | 5’TTCAGAGGATGCTCTAGGGAAGA 3’ | 5’CTGTGGAGTAAGTCCTGTGCC 3’ |
| *Kdm5d* transgene | RT-qPCR | 5’GTGGAGCCACCCCCAGTTC 3’ | 5’GAAAGTCGTCAGATCCTGGCTTC 3’ |
| *Kdm5d* (total) | RT-qPCR | 5’GCAGCATTGAGGAGGATGTG 3’ | 5’CCCAGTGCAGGTAGTTAATGG 3’ |

[a] Exons 11 and 12 are underlined. loxP sites are shown in bold. Intronic sequences are shown in lowercase.
[b] Gene names are italicized.
